# Gradual ontogenetic shifts in the mobility and space use of long-lived migratory greater flamingos

**DOI:** 10.1101/2025.06.16.659883

**Authors:** Lina Lopez-Ricaurte, Wouter M.G. Vansteelant, Arnaud Antoine, Olivier Duriez, Frédéric Jiguet, Sergio Nissardi, Davide Scridel, Lorenzo Serra, Stéphan Tillo, Arnaud Béchet, Jocelyn Champagnon

## Abstract

1. How early-life exploration shapes the adult annual cycle routines of migratory animals remains challenging to study, especially in long-lived species. Delayed recruitment often observed in long-lived migratory birds suggests that the first breeding attempt may be constrained by a protracted learning process in which individuals develop their annual itinerary, which may involve extensive exploration of the winter and breeding sites, potentially causing inexperienced young to wander beyond traditional population areas.
2. Using greater flamingo *Phoenicopterus roseus* GPS tracking data from 83 individuals tagged as nestlings in the Mediterranean, we analyzed ontogenetic changes in mobility and space use for up to 8 years of age. We segmented autumn-winter and spring-summer tracks (n = 223 device-bird-year combinations) into staging events (n = 1914). We then computed seasonal staging metrics and analysed how they changed with age. We map the traditional congregation sites of flamingos to investigate age-related changes in space use between males and females.
3. Flamingos were more exploratory during their early years, gradually transitioning to a more sedentary lifestyle as they grew older. With each additional year of age, the number of staging events decreased by 16%, while the average duration of staging events increased by 8 days.
4. Birds remained faithful to the non-breeding staging sites where they spent most of their time during the first year of life. Over time, they returned progressively closer to their natal sites, even after years of exploration at non-traditional locations. Breeding was rare (8%) and occurred at a relatively late age (≥5 years), highlighting an extended exploratory phase, a pattern observed in other long-lived species.
5. Older birds were more likely to use traditional sites and spent more time at these sites, particularly after age 5. Our study reveals the individual exploration process underlying natal and non-breeding dispersal patterns previously laid bare by ring-resighting studies and shows that flamingos retain long-term memory of sites learned in early life.

## Introduction

Across the animal kingdom, both invertebrates and vertebrates engage in exploratory movements, a behaviour observed across various body sizes and ecological contexts (Züst et al. 2023 and references therein). Generally, individuals gain knowledge through exploration, defined as the acquisition of information by an animal while navigating through unfamiliar environments (Verbeek et al. 1994). Exploration likely involves a trade-off between the risks of venturing into unfamiliar areas (which may include poorly protected areas) and the benefits of gaining experience (e.g., to locate food, shelter, or suitable habitats later in life, Smithson & Johnston, 1999; Brightsmith et al., 2021). In highly mobile species, how a young individual explores and learns during its early life is crucial for its survival and reproductive success (African cichlid *Simochromis pleurospilus*, Kotrschal & Taborsky, 2010); painted turtle *Chrysemys picta*, (Roth & Krochmal, 2015) In mammals, exploratory movements typically peak during early life and may be precursors to successful dispersal and ultimately responsible for gene flow (Debeffe et al., 2013).

In long-lived migratory birds, early-life exploration is particularly complex and typically involves multiple stages. Young birds must learn outward migration routes, including stopover sites, and establish non-breeding locations during their first year of life. This is often followed by a wandering phase that may span multiple years, during which individuals expand their knowledge of the world. Eventually, they learn the return migration routes and settle at breeding sites, completing the natal dispersal process (Green et al., 1989; Betts et al., 2008). Once recruited, adult breeders tend to be faithful to their breeding and non-breeding sites in consecutive years (Newton, 2012). Nevertheless, even experienced breeders may disperse to alternative breeding or non-breeding sites, and such events are likely informed by early-life experience too (Vickers et al., 2023). As such, early-life exploration underpins many important ecological and evolutionary processes, from natal dispersal, recruitment, and individual fitness (Nathan et al., 2008) to metapopulation dynamics (Johst et al., 2002) and species distributions (Greenwood, 1980; Greenwood & Harvey, 1982).

Early-life processes such as natal dispersal have perhaps been best studied in short-lived bird species such as songbirds (Weatherhead & Forbes, 1994), which have relatively short parental care followed by a brief period of early-life exploration before settling to breed in their second calendar year (i.e. one year old) (Baker, 1993). By contrast, long-lived species have a protracted early-life period spanning several years, with recruitment often delayed well beyond sexual maturity and considerable inter-individual variation in the age of first breeding. For example, common terns *Sterna Hirundo* typically return to the colony at age 2, but first breed between 3 and 5 years (Szostek & Becker, 2012). In more extreme cases, bearded vultures *Gypaetus barbatus* return to their breeding areas around age 7 and recruit at ∼10 years, while wandering albatrosses *Diomedea exulans* return at 5–7 years and recruit between 7 and 15 years (López-López et al., 2013; Weimerskirch et al., 2005). And it is intriguing that despite such different early-life trajectories, the norm for most species is (i) to return near their natal sites to breed (Coulson, 2016), (ii) to be highly faithful to breeding (Newton & Marquiss, 1982) as well as (iii) non-breeding sites (Lourenço et al., 2016). However, the intricacies and functionality of early-life exploration remain obfuscated by the difficulty of tracking long-lived species from fledging to adulthood. In particular, understanding how and when young birds establish adult routines and select breeding and non-breeding sites—often maintained for life—remains a challenging frontier. Bridging these knowledge gaps is essential to understand how early experiences shape the long-term ecological and evolutionary trajectories of these species.

Limited studies on early-life intricacies in long-lived species, revealed by few tracking studies, pointed out considerable individual variation of migratory behaviour within individuals of the same population. This individual variation has been partially attributed to a high degree of plasticity during early life in response to environmental and social dimensions (Battley et al., 2020; Verhoeven et al., 2022), a phenomenon referred to as developmental plasticity (Piersma & Drent, 2003). Such developmental plasticity is particularly evident in flying and swimming organisms, with the establishment of individual non-breeding sites being strongly influenced by stochastic wind conditions experienced by first-time migrant birds (Vansteelant *et al*. 2017) and establishment of migratory routines being strongly influenced by interannual variation in hatching drift patterns in sea turtles (Scott et al., 2014). Additionally, social learning, potentially acquired from related (Brent goose *Branta bernicla*, Harrison et al., 2010) or unrelated conspecific (lesser spotted eagle *Clanga pomarina*, Meyburg et al., 2017), but also from co-occurring species (whooping cranes *Grus americana*, Teitelbaum et al., 2019; northern bald ibis *Geronticus eremita,* Piersma, 2024), may influence both the choice and the propensity of youngsters to switch staging areas from year to year. Typically, adults are more faithful to a single staging site in consecutive years, in contrast to younger birds, which actively explore and sample the environment. In Eurasian spoonbills *Platalea leucorodia* on their first southward migration from the Netherlands, the higher probability of young individuals switching wintering sites in consecutive years was attributed to early life exploration (Lok et al., 2011). Sex is an important factor mediating inter-individual variation in early life processes. If males are the territory-establishing sex, female-biased natal dispersal is often observed (in fish, Taylor et al., 2003; mammals, Lawson Handley & Perrin, 2007; butterflies, Stevens et al., 2011).

Here, we use a longitudinal GPS tracking data set to investigate ontogenetic shifts in the mobility of greater flamingos *Phoenicopterus roseus* originating from different Mediterranean breeding colonies, from fledging up to 8 years of age. Previous work using ring resightings revealed that greater flamingos born in the Camargue (South of France) showed high site fidelity to wintering sites from the first year onward (Barbraud et al., 2003; Sanz-Aguilar et al., 2012). Balkız *et al*. (2010) revealed that most individuals born in the western Mediterranean (i.e., the Camargue, Fuente de Piedra, Spain, and Molentargius, Sardinia) do not make their first breeding attempt in their natal colony. Instead, they dispersed to other colonies (natal dispersal) within the Mediterranean basin before returning to their natal site to breed later in life, a behaviour observed in other colonial breeding species (Pakanen et al., 2022; Péron et al., 2010). How early life exploration mediates recruitment deserves further investigation.

Following what we know about early-life exploration in long-lived species, we hypothesize that younger birds exhibit a linear decrease in exploratory behaviour throughout the years. As they age, this behaviour is expected to transition into more seasonal movements between a smaller number of staging sites. Consequently, we predict: (1) a rapid decrease in the number of staging events and an increase in the mean duration of staging events over time; and (2) a significantly higher number of staging events with shorter durations in smaller females compared to larger males, due to weaker competition for securing high-quality territories among the former. In species with sexual size dimorphism, moving is theoretically less costly for the smaller sex (Li & Kokko, 2019), thus we predict (3) that larger males will be more sedentary than smaller females. Based on previous studies with ringing-resighting data, we predict that (4) greater flamingos will remain highly faithful to staging sites where they spent most time during their first winter (Sanz-Aguilar et al., 2012; Barbraud et al., 2003; Green et al., 1989) and that summer staging sites will be located progressively closer to the natal site with age in line with Balkız et al. (2010).

We tested non-exclusive hypotheses regarding how age, sex, and season affect mobility during early-life. If dominance and individual quality increase with age, the dominance hypothesis (Townshend, 1985) predicts that smaller, younger individuals are displaced from the best habitats but gradually move to better sites, in terms of fitness, as they age. Assuming that greater flamingos travel in flocks consisting of inexperienced birds (Johnson and Cézilly 2007), we predict that younger individuals are more likely to spend time outside of population areas or sites where large aggregations of greater flamingos are known to breed or spend the non-breeding season (hereafter traditional sites) during both breeding and non-breeding seasons. As they age and become more dominant and capable of coping with conspecific competitors, they will gradually spend more time in traditional sites, and males will stay longer relative to females. Alternatively, if greater flamingos travel in mixed-age flocks of conspecifics, social learning might ensure the same traditional sites are being used at all ages, even during the first outward migration. Furthermore, first-time breeders (low-competitive individuals) are expected to settle first into low-quality, low-density colonies to acquire breeding skills (the breeding-skill hypothesis; Ens et al., 1995). In subsequent years, after gaining reproductive competence, individuals may return to their natal colony (a phenomenon referred to as delayed natal philopatry; Balkız et al. 2010). We predict that a higher proportion of natal dispersers will make their first breeding attempt elsewhere but will later return to breed at their natal site.

## Materials and methods

### Study system

The greater flamingo is a long-lived colonial waterbird with a body mass dimorphism, with males being ca. 20% heavier than females. In the western Palearctic, greater flamingos regularly migrate between breeding and non-breeding areas in the Mediterranean basin along the East Atlantic or the Central Mediterranean migration flyways, with a minority of individuals residing close to their natal area. Greater flamingos congregate at well-known traditional sites in the Mediterranean during both breeding and non-breeding seasons where colonies of > 20.000 breeding pairs are known. Previous studies have demonstrated that most of the greater flamingos in Camargue are 4-6 years old when they first breed (Johnson & Cézilly, 2007).

### Tagging and tracking

From 2015 to 2024, 101 greater flamingo (15 females, 34 males, and 52 of unknown sex) were fitted with GPS loggers just before the fledging (Table S1). This was achieved by driving part of the chick crèche into a corral during the annual ringing operations of the species (see Johnson & Cézilly, 2007 for details). Captures were made in one colony in France and three different colonies in Italy: (1) Aigues-Mortes salt pans (north-western Mediterranean Sea, 43.55 ° N, 4.18 ° E), (2) Molentargius salt pans in Sardinia (central Mediterranean Sea, 39.23 ° N, 9.12 ° E), (3) Comacchio lagoon and salt pans (north-western Adriatic Sea, 44.64 ° N, 12.21 ° E) and (4) Margherita di Savoia salt pans (Adriatic Sea near the Gulf of Manfredonia, 41.39 ° N, 16.01 ° E) (Figure 1A). Tagged birds were ringed and measured and for each bird born in Italy, molecular sexing was carried out (Scridel et al., 2023). Permits for tagging and tracking in Italy were under authorization of Law 157/1992 [Art. 4(1) and Art. 7(5)]. Ringing and tagging in France were conducted under regulatory authorizations from the National Ringing Centre (CRBPO) as part of programs licensed under permits PP1190 and PP405. (Appendix 1).

**Figure 1.**
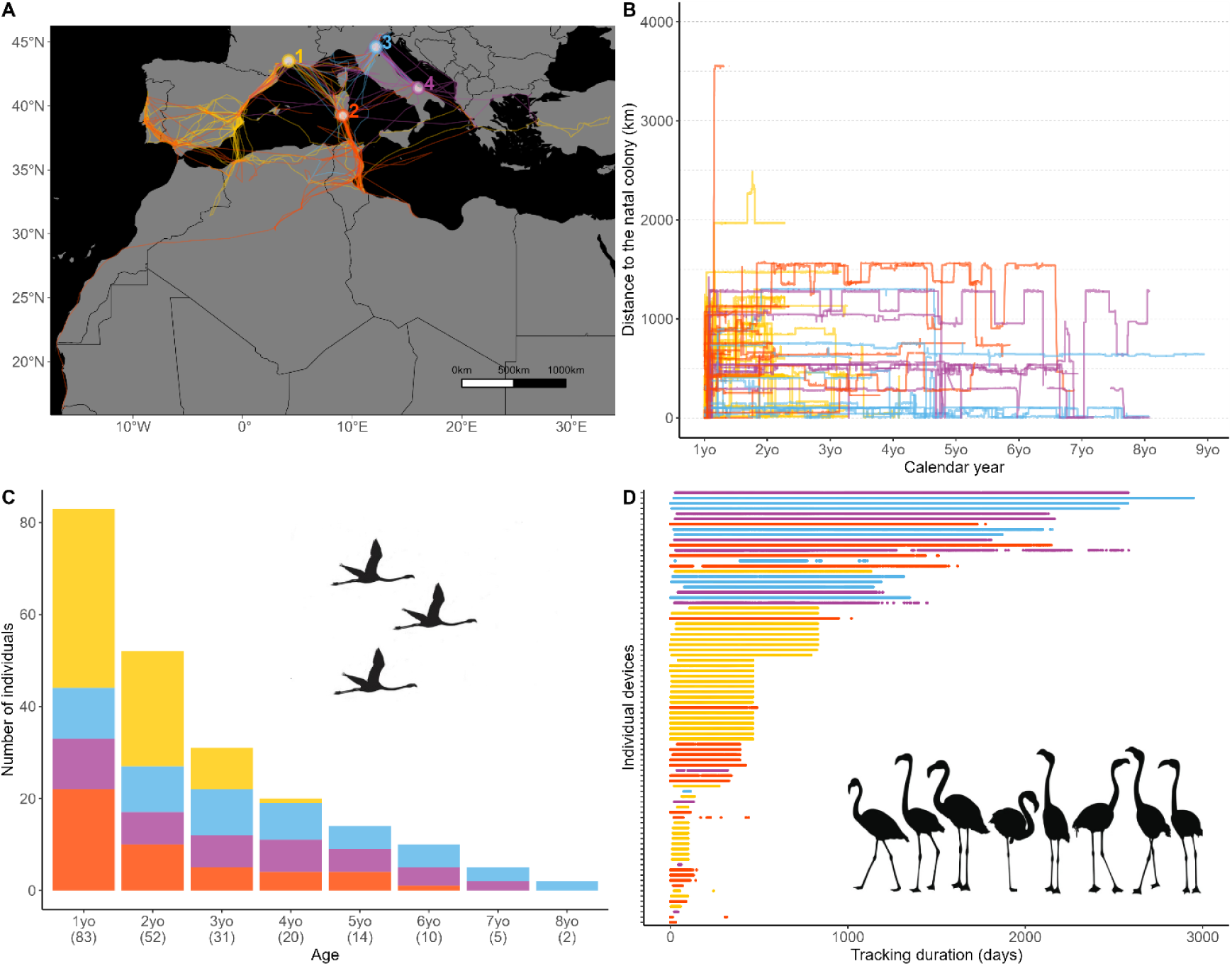
GPS-tracking data used in this study from 84 greater flamingos (*Phoenicopterus roseus*) tagged as fledglings in Southern France and Italy. A. Study breeding locations and full tracks of all birds. Lines and bars are coloured according to the colony: (1) Aigues-Mortes (43.55° N, 4.18° E) in yellow, (2) Molentargius (39.23° N, 9.12° E) in orange red, (3) Comacchio (44.64° N, 12.21° E) in blue and (4) Margherita di Savoia (41.39° N, 16.01° E) in purple. B. Timing of movements shown as the increasing or decreasing distance to the natal colony throughout the study period. Number of individuals tracked each calendar year by the colony. The numbers of unique individuals tracked each year are indicated in parentheses, with individuals tracked over consecutive years appearing in multiple calendar years D. Tracking duration in days since the onset of post-fledging dispersal of each individual by colony. The gaps indicate periods in which no data were recorded.

For our analyses, we included 83 individuals, 223 device-bird-year events and 391 seasonal events, from which 217 were classified as autumn-winter and 174 as spring-summer (unique combinations of individual, bird-year, and season). For details on excluded birds, see Appendix 2.

### Resampling and track annotation

Due to the slightly different sampling frequencies, all data were resampled to a 1 h interval, allowing deviations up to 20 min. By resampling, we avoided bias in movement metrics that may result from variability in sampling rates (Shamoun-Baranes et al., 2016; Vansteelant et al., 2015). The resampled data set comprised c. 682,000 locations. We filtered and excluded unrealistic average ground speeds (i.e. when the speed between two consecutive locations was over 30 m/s). Movement metrics (distance, duration and speed) were calculated from each resampled fix to the next, applying the Haversine Distance formula with the deg.dist function in the R package fossil (Vavrek, 2012).

To annotate GPS locations within traditional sites, we used the polygons of Important Bird and Biodiversity Areas (IBAs) designated specifically for greater flamingos (BirdLife International, 2024), which captures 92 % of the traditional sites known to the flamingo specialist involved in this study. For the 8 % known traditional sites that were not designated as flamingo IBA, we created custom polygons using the map of flamingo breeding colony locations updated during the 7th Greater Flamingo Workshop in the Mediterranean and West Africa, 2024 (hereafter the 7th GFW) (Table S2).

### Track segmentation

Individual tracks were segmented into consecutive age classes: 1yo (1-year-old), 2yo, 3yo, 4yo, 5yo, 6yo, 7yo, 8yo. To do this, we used a period of 365 days, starting from the post-fledging dispersal date of each individual: i.e. the departure of a juvenile from its natal area (post-fledging dispersal) as the day when the bird flew over 100 km away from its natal colony and did not return for the following 15 days (Scridel et al. 2023). We verified these dates by examining each individual’s GPS tracks. To identify staging events (i.e., resting or foraging periods) from directed movements (i.e., travel periods), we used an hourly speed threshold of 5 km/h, which was defined by identifying a break in the histogram of ground speeds (Figure S1). Then we segmented the tracks into bouts of continuous staging and travel events. As we were interested in staging behavior, we excluded all travel events. Finally, we labeled staging events as inside or outside traditional sites and calculated the duration of each staging event (in days).

### Staging metrics

We used 673,015 staging events of greater flamingos to differentiate short vs long visits. We classified visits as short when the individual just passed over and long visits when an individual used a site for several days in a row (at least 3 days in which a bird roosts less than 10 km apart). From each long visit, we determine the median longitude and latitude, the arrival and departure dates, and the duration. We repeated our data analyses with different temporal thresholds to qualify a long stay (that is, 3, 7 and 10 days) and found no difference in the results (Figure S2). For each individual-age-season event, we identified the main staging site as the most used site during summer-spring and autumn-winter, i.e., as the site where the birds spend most of the time for a given season. We then calculate the distance from the previous main sites in consecutive years in each autumn-winter and spring-summer period as the straight-line distance (i.e. the shortest orthodromic route). Finally, we calculate the straight-line distance to the natal colony of those main sites.

### Reproductive status

In the greater flamingo the incubation period lasts c. 29 days (range: 26-32, Johnson and Cézilly 2007). About two days after the egg is laid, one parent leaves to feed, usually for 1 to 4 days, then returns to swap the incubation shift with the other. We extracted the breeding status of our tracked birds by visually inspecting the time series of latitudes and longitudes. We filtered the months from April to June (i.e., the period of the first clutch of the greater flamingo in our study breeding colonies) from age 3 onward (the earliest observed age of first reproduction in the species (Tavecchia et al., 2001), resulting in 66 combinations animal-age. We then mapped these tracks with the locations of breeding colonies within the Mediterranean basin (N= 71, source 7th GFW). We extracted individuals that were stationary within a 1 km buffer of a known flamingo colony during the breeding period. For each combination of individual and age, we searched for 2- to 4-day time windows and classified birds as ‘potential breeders’ or ‘failed breeders’. A bird was labelled a ‘potential breeder’ if it remained at the colony site for at least 4-time windows. It was labeled a ‘failed breeder’ if it showed less than 2 incubation shifts before leaving the colony or spent long periods in the colony (more than 9 days, likely due to molt initiation after incubation failure, Johnson and Cézilly 2007). Finally, we classified as ‘visits’ the instances in which an individual came inside the 1 km buffer for 1-4 days and then left the colony without returning.

### Statistical analyses

To identify whether there is a significant change in staging metrics (i.e. the number of staging events, the average staging duration (in days), the distance of main staging sites to the natal colony (km) and the distance from the previous main staging sites (km), between seasons, ages and sex, we ran generalized linear mixed models (GLMMs). For the number of staging events (response variable), we fitted a Poisson distribution with a log link function. For the average staging duration (response variable), we fitted a Linear Mixed Effect Model (LMM) with Gaussian error distribution. We included bird identity, year, and colony as random intercepts. We used GLMM with a Gamma distribution (proper for positive continuous variables, Zuur, 2009) and a log link function for the analyses of the distance of main sites to the natal colony and for the distance from previous main sites. To reduce model complexity, we removed random effects with a variance of zero or close to zero (i.e., year and colony). As fixed effects, we included age (continuous variable, centred and standardized, Schielzeth, 2010), season (autumn-winter: September-February and spring-summer: March-August) and their interaction. Next, we tested whether birds increased the use of traditional flamingo sites with age by fitting an LMM for the mean duration of the segment (response variable). As fixed effects, we included traditional sites (two-level categorical variables: inside, outside) and season. Finally, we repeat all the analyzes using a subset of the dataset that included only individuals for which sex was determined, incorporating sex as fixed effects along with its interaction with age. We excluded older individuals (i.e., year-5 and older) due to unbalanced sex ratios (Table S3). We examined strong support for an effect whenever 95% confidence intervals of the coefficient excluded zero (Nakagawa & Cuthill, 2007). The proportion of variance explained by the fixed effects (Rmarginal) and by both fixed and random effects (Rconditional) was assessed using the methods in Nakagawa & Schielzeth (2013).

To assess the relationship between age and the response variables, we modelled age using four different functional relationships (linear, quadratic, exponential and cubic; see Appendix 3 for details). For all models, we checked for potential residual overdispersion of residuals using the R package DHARMa (Hartig, 2020) and visually inspected the residual plots. We use Akaike’s information criterion (AIC; Burnham & Anderson, 2002) to rank all sets of models resulting from each response variable and select the best model (that is, with the lowest AIC) for further analyzes (Table S4). When the difference among the top models was ≤2 units, we retained the most parsimonious model (Burnham & Anderson, 2002). Once the functional relationship of age for each response variable was selected, we used the ‘dredge’ function in the MuMIn package R (Barto, 2025), which uses Akaike’s information criterion (AIC) to rank all possible subsets of reduced models from each best model. All data processing and analyzes were performed using the R Language for Statistical Programming, version 4.2.2 (R Core Developmental Team, 2022).

## Results

Greater flamingos left their natal area between August 6 and December 16 (median ± SD: September 9 ± 23 days; n = 83 post-fledging dispersal tracks; Figure S3A). Post-fledging departure timing differed significantly among colonies, with Molentargius birds departing earlier than those of other colonies (F₃,₈₀ = 9.79, P < 0.001; Figure S3B). The staging sites of the tracked birds during the autumn-winter (non-breeding) and spring-summer (breeding) seasons were located on a broad front, from the Portuguese Atlantic coast to the north-western Turkish coast (Figure 1A). Only one 1 yo male flamingo born in Molentargius (Italy) moved out of the Mediterranean region, crossing the Sahara Desert from Tunisia to the Atlantic coast of Morocco, and then following a coastal route until its final destination at the Mauritanian-Senegalese border.

### Age effect on staging metrics

Age-related changes in the number of staging events between seasons were best described by a negative linear relationship (Table S4, Table S5). The most parsimonious model included age and season (Table S6), indicating that younger birds visited many more sites for shorter periods compared to older ones (β = -0.17 ± 0.04, CIs: -0.25, -0.10; Figure 2A, Table S7). All age classes exhibited a greater number of staging sites during autumn-winter/September-February (mean ± SD: 5.4 ± 3.42) than spring-summer/March-August (4.6 ± 3.54). The interaction between age and sex was significant (β = -0.32 ± 0.11, CIs: -0.54, -0.11), indicating that older males visit fewer sites than older females (Table S8, S9). The average staging duration increased linearly as greater flamingos grew older by 8 days per year of age (β = 7.99 ± 3.49, CIs: 1.12, 14.85; Figure 2B, Table S7). The most parsimonious model retained the interaction between age and season (Table S6), indicating that seasonal differences became progressively higher with age, with longer staging events during spring-summer (60.62 ± 53.80 days) than autumn-winter (39.65 ± 37.90 days). The interaction between age and sex was significant (β =18.87 ± 7.59, CIs: 3.90, 33.83), indicating that the duration of the staging was longer for older males than for older females (Table S9).

**Figure 2.**
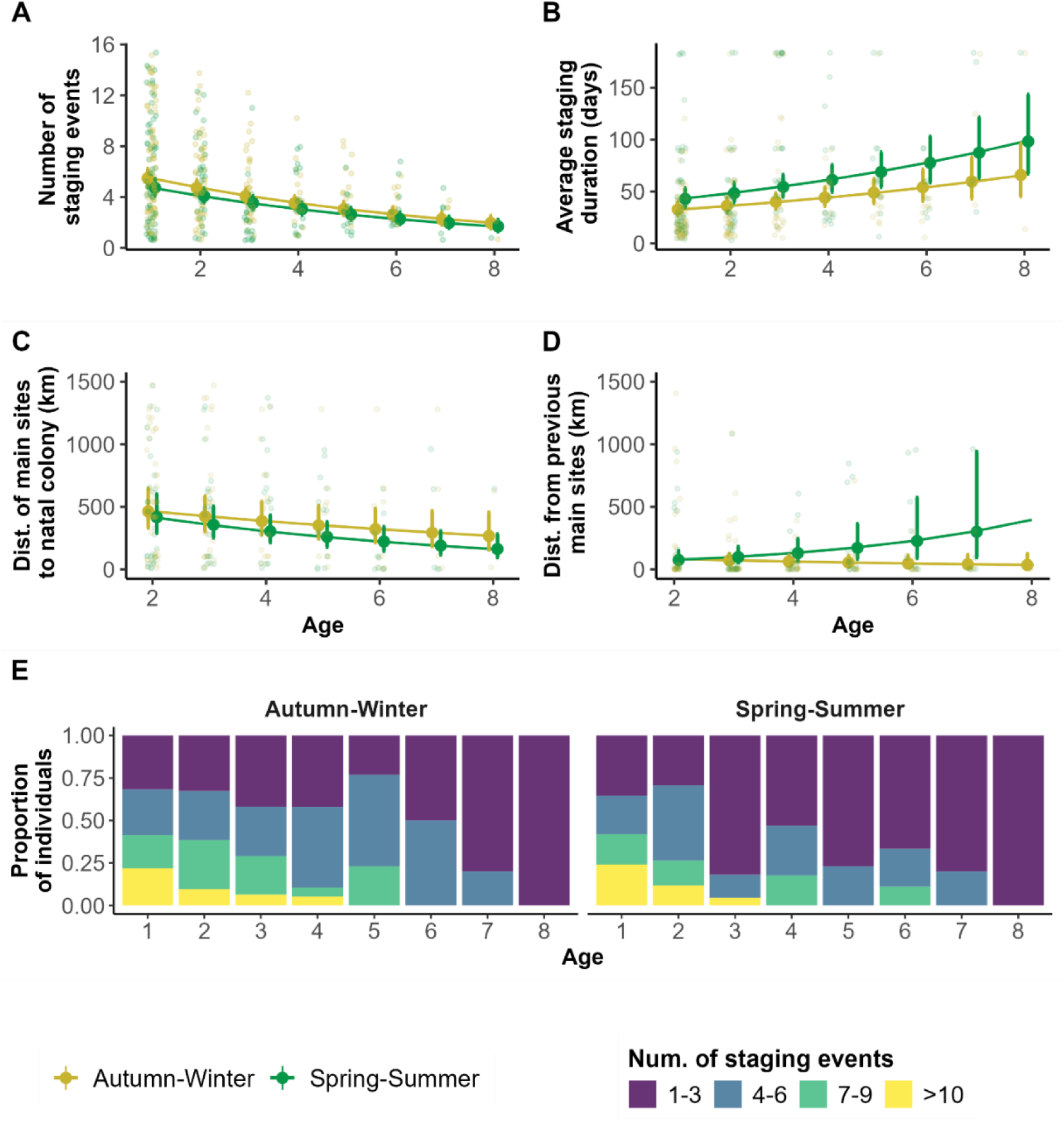
Ontogenetic changes in mobility of greater flamingos during autumn-winter/non-breeding (yellow) and spring-summer/breeding season (green). (A) The number of staging events as a function of age and season. We classify staging events as periods >3 days in which a bird roosts less than 10 km apart (n=1914 staging events). (B) The average staging duration is in days. (C) The distance of the main staging sites (i.e. longest duration) to the natal colony. (D) The distances from the main staging site to the previous main staging site in consecutive years in each breeding and non-breeding season. (E) Proportion of birds with different numbers of staging events over time. The line represents the model estimates ± the 95% confidence interval.

### Site fidelity and philopatry

We found a negative linear relationship between age and the distance between the main staging sites and the natal colony (Figure 2C). The most parsimonious model included age and season (Table S6), indicating a linear decrease with age (β = -0.19 ± 0.05, CIs: -0.30, -0.08; Table S7) and an overall shorter distance to the natal colony during spring-summer than during autumn-winter (β = -0.21 ± 0.09, CIs: -0.39, -0.03, Table S7). Furthermore, a linear relationship best described age-related changes in the distance from the previous main staging site during the autumn-winter and spring-summer sites (Table S4). The most parsimonious model retained the interaction between age and season (β = 0.67 ± 0.26, CIs: 0.15, 1.18, Table S5, Table S6). Specifically, the distance between sites used in consecutive autumn-winter seasons became very flat and low from the 2nd year of age onwards, indicating high fidelity to non-breeding areas. During spring-summer, the distance between consecutive staging sites increased significantly with age, with no evidence of sex differences in this exploratory behavior (Table S9).

### Traditional sites and experience

Male and female flamingos frequently used traditional and non-traditional sites during their early life (Figure 3B). On the contrary, the majority of experienced individuals barely stop outside traditional sites (Figure 3C). The most parsimonious model retained the interaction between age and traditional sites (Table S10). The time spent within traditional sites showed a significant exponential relationship (β = -3.69 ± 0.88, CIs: -5.41, -1.96, Table S7), with a notable decrease, reaching near zero values from the age of 5 years onwards (Figure 3D). We did not find evidence of season or sex differences (Table S8, S9).

**Figure 3.**
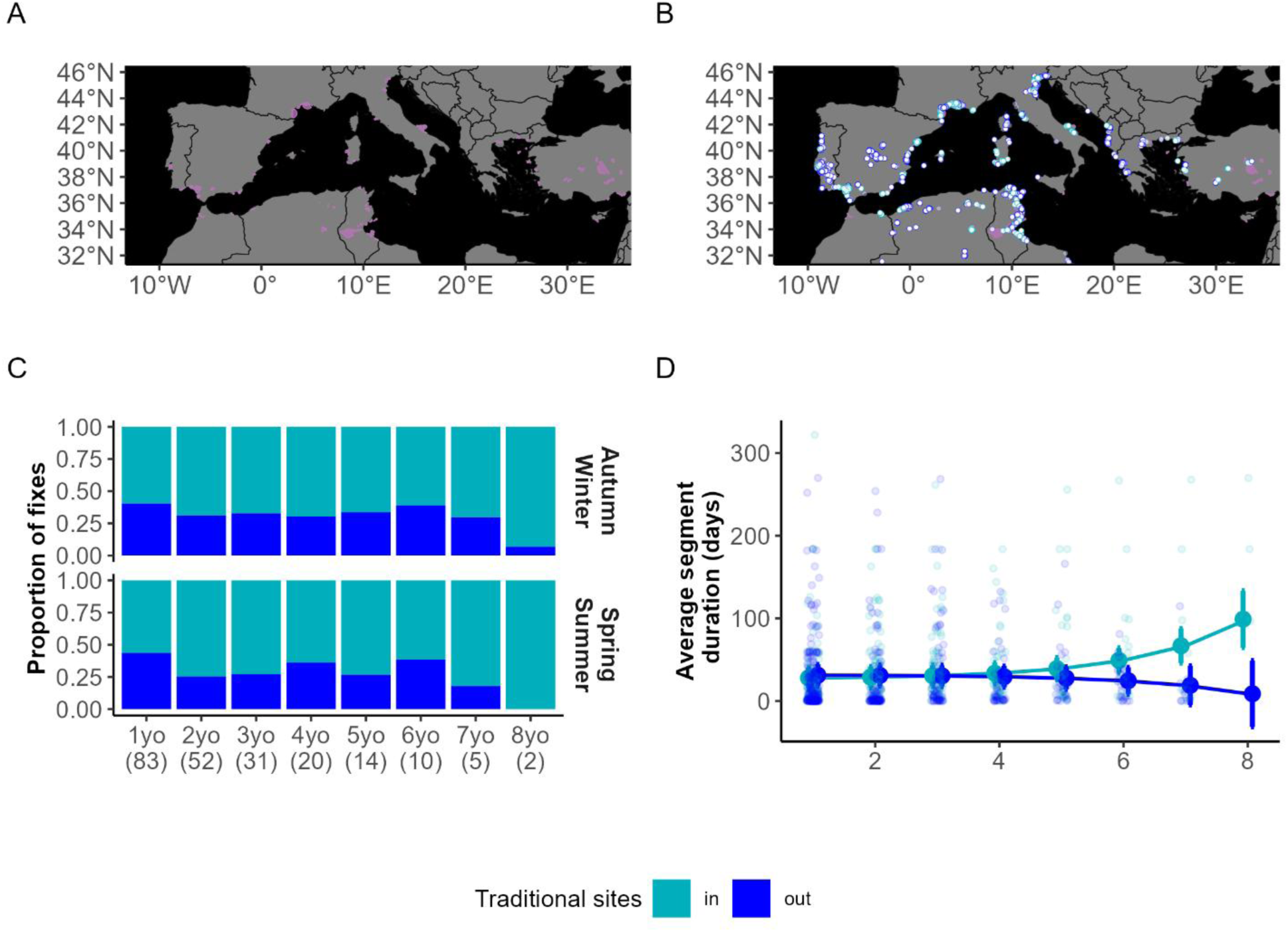
Flamingos’ traditional congregation sites. **(A)** Map of traditional sites (pink areas) by greater flamingos year-round. We combined IBAs designated specifically for flamingos with information from the map of the breeding colonies produced by experts during the 7th GFW. **(B)** Location of fixes in and outside traditional sites. Aquamarine colour represents fixes inside and navy blue outside traditional sites. **(C)** Age differences in the proportion of fixes inside and outside traditional sites. **(D)** Average time spent in and out of traditional sites over time. The line represents the model estimates ± the 95% confidence interval.

### Reproductive status

26 individuals were found to stage within 1 km buffer of a known flamingo colony during the breeding period. Of these staging events in a colony, 32 % (n=21/66) were classified as visits and 8 % (n =5/66) were classified as potential breeders or failed breeders (Figure S4). The average age of the onset of potential breeding behavior was 5 years (range: 4 to 7 years). Finally, of the potential breeders or failed attempts, 3 were in the natal colony, while 2 switched to another colony. One of these birds attempted to breed in two consecutive years, at 6 and 7 years old, and consistently returned to the same breeding colony.

## Discussion

Using advanced tracking technology, we verified the patterns identified from ring-resighting data in previous studies, while for the first time revealing the flexible early-life movement process that generates repeatable site use routines of long-lived greater flamingos. These conclusions add evidence of the remarkable capacity of birds to learn, remember and navigate back to previously learned sites, and the key role of early-life exploration to build knowledge that can help long-lived species thrive in fluctuating environmental conditions.

As expected, the mobility and space used by the greater flamingos gradually changed as the birds grew, with the older birds restricting their activities to fewer sites for longer periods. These results are in line with predictions derived from previous work that juveniles are more likely to explore new areas than adults (Morand-Ferron et al., 2011; Tombre et al., 2019; Wolfson et al., 2019). Explorative behaviour by juveniles can be attributed to fitness consequences and experience. In long-lived species, the age of first reproduction may be delayed up to several years, and in our dataset, we found that the earliest age of first potential breeding was 5 yo (cf. Johnson & Cézilly, 2007). This relatively late age of first breeding in greater flamingos allows for a more prolonged exploratory phase that might be part of cognitive processes about resources and stressors (e.g., mates, predators, food acquisition). In this sense, there are likely to be fitness and survival benefits associated with the early-life exploration of alternative breeding and non-breeding sites, including non-traditional flamingo sites (Stearns, 1992; Lewis et al., 2021). More specifically, early-life exploration probably allows birds to build a larger mental map of potential breeding and non-breeding areas and may enable flamingos to cope with annual fluctuations in breeding conditions later in life. Indeed, traditional colonies can be abandoned for up to several years if conditions are not permissive for breeding (e.g., drought), in which case, experienced adults may draw on early-life experience to attempt breeding elsewhere, rather than skipping breeding altogether. Contrary to our prediction, greater flamingos did not show sex differences in most staging metrics, in line with previous studies on natal dispersal for this species (Nager et al., 1996; Barbraud et al., 2003). However, our models did capture an effect of the interaction between age and sex in the number and duration of staging events. Older males visited significantly fewer sites for longer periods than older females. Although small sample sizes urge cautious interpretation of these results, the higher mobility of older females aligns with the general pattern observed in bird and mammal species, where females and juveniles often disperse more than males (Greenwood, 1980).

Our longitudinal tracking data suggest a long-term memory of traditional breeding and non-breeding sites in greater flamingos (Figures 2C and 2D). They have the cognitive ability to return to previously learned non-breeding destinations (Sanz-Aguilar et al., 2012; Barbraud et al., 2003; Green et al., 1989). In addition, they show a high degree of natal philopatry, with individuals moving progressively closer to their natal area over successive summers, even many years after birth. This occurs after having built up much experience at alternative traditional sites and non-traditional sites. Some cases of extreme early-life exploration and long-term memory of natal areas have been observed for European honey buzzards *Pernis apivorus*, which may not return to their natal sites until 3-4 years after their first migration, and with some individuals undertaking wide-ranging exploratory movements for several summers before finally returning to natal sites to breed (Mirski et al., in review). In line with previous studies, we found that 3 probable breeding attempts were made in the natal colony while 2 were in another colony, confirming philopatry or delayed philopatry of this species (Balkız et al. 2010). However, the latter study relied on a data set with many more older individuals and for a longer period (20-year capture-mark-resighting dataset) and as flamingos can live up to 40 years in the wild (Johnson and Cézilly, 2007), our 8-year study captures only part of the dispersal capacity and flexibility of the species.

Already from the first year onward, tracked flamingos discovered species’ traditional sites, suggesting that naive juveniles may learn from experienced elders (social learning) to reach such congregation sites. In species such as swans, geese, and cranes, young birds migrating in family units are known to learn migration routes from their parents. Our findings add to a growing body of evidence that social learning of migration as well as breeding and non-breeding destinations is likely a far more widespread phenomenon, including species in which parental care ends before the first migrations, such as the greater flamingo (Aikens et al., 2024, Newton 2024, Lok et al., 2025). In addition, the time spent at non-traditional sites decreased exponentially and finally reached near-zero levels from the age of 5 years onwards. This is in agreement with predictions from the dominance hypothesis, whereby dominant high-quality experienced flamingos occupy traditional sites, while inexperienced flamingos are pushed out to non-traditional sites (Marques et al., 2010; Lok et al., 2011).

As expected and in accordance with previous studies (Green et al., 1989; Barbraud et al., 2003; Sanz-Aguilar et al., 2012), most flamingos discover their primary autumn-winter sites before year 2 and remain faithful to these sites thereafter. This suggests that flamingos derive significant benefits from returning to familiar sites, often selected during their first non-breeding season. These benefits are likely related to familiarity with resource availability, predator presence, and other potential dangers, which can enhance individual fitness (Lewis et al., 2021; Riotte-Lambert & Weimerskirch, 2013). During spring-summer the distance between main staging sites increased linearly across consecutive years. At first, this seems like a counterintuitive result. Why are we not seeing a decreasing distance as birds settle at specific sites? We hypothesize that despite being physiologically mature at 3 years of age, they still need to refine their reproductive skills (e.g. group displays, Johnson and Cézilly, 2007) and acquire public information by visiting various colonies. This learning and social process extends beyond the age of 8 years as observed in other long-lived species (Sergio et al., 2014; Campioni et al., 2017). Alternatively, they may have sufficient skills to breed, but are excluded due to density-dependent intraspecific competition for nesting sites and mates, prompting them to move elsewhere in search of less competitive colonies or delayed reproduction (queue for vacancies, Forbes & Kaiser, 1994; Ens et al., 1995; Kokko & Johnstone, 1999). Furthermore, when looking at the individual level, we observed great variation between individuals (Figure S5). We suspect that the observed increase in the distance between main staging sites is driven by a few individuals tracked over a long period who dispersed exceptionally long distances, strongly shaping the overall trend in older age classes.

Using GPS technology, we tested hypotheses previously explored with ring-resighting data, overcoming limitations such as detection probabilities and observer bias. Our findings highlight the processes of early-life exploration as a key driver of site fidelity in both breeding and non-breeding contexts for long-lived species. In our study system, greater flamingos demonstrate “flexible routines”, where early-life exploration helps establish preferences for breeding and non-breeding sites, while building experience and familiarity with sites to which they may disperse when environmental conditions necessitate. Such flexibility is likely advantageous in the face of large inter-annual fluctuations in Mediterranean precipitation. An important avenue for future research is to explore whether early-life experiences and memories have lasting consequences throughout an individual’s life. For instance, do bolder juveniles have higher survival rates or greater breeding success compared to more cautious individuals? How do experienced individuals navigate the balance between personal knowledge and social information? Longitudinal studies tracking individuals from early life to adulthood will be essential to answer these questions and provide deeper insight into how early-life decision-making shapes adult behaviours and routines.

