## Supplementary material for "Gradual ontogenetic shifts in the mobility and space use of long-lived migratory greater flamingos": Suplementary Information

**Appendix 1**

Juvenile greater flamingos were tagged with 5 models of 25 g, solar GSM-GPS devices from different manufacturers (OMNI 2G, Interrex, Poland; Ornitrack-25, Ornitela, Lithuania; SAKER-H and SAKER-L, Ecotone, Poland; GsmTag-U9, Milsar, Cyprus), attached as backpacks with a Teflon harness. The Interrex devices were mounted on the leg, specifically on one of the tarsi. Total backpack weight (i.e. device plus harness) was below 3% of mean body mass, which is within the accepted standards for animal welfare in research (Phillips et al., 2003; Barron et al., 2010). All devices recorded geographical locations day and night at intervals ranging from 1 to 2 hours. Birds in Italy were ringed and tagged by the Italian Institute for Environmental Protection and Research (ISPRA) from 2015 to 2017 and in France by Tour du Valat from 2021 to 2023 (Figure 1, Table 1s).

**Appendix 2**

Of the birds excluded (18 out of 101), 7 were confirmed dead (corpses retrieved) before providing useful data, that is: 6 died before post-fledging dispersal. We also retrieved the carcass of 1 flamingo on the northwestern coast of Sidi Ghiles, Algeria, who died after embarking on a first non-stop Mediterranean crossing, one month after being tagged in France. 5 tags stopped providing GPS data shortly after deployment (minimum = 18 days, maximum = 70 days) before the birds left the natal colony. 1 bird tagged in Margherita di Savoia did not yield any data, and 5 resident birds that remained within the system of wetlands near their natal area (i.e., maximum beeline distance <70 km staging for more than four months, min = 136 days, max = 636 days) were excluded from the analyses, as their year-round movements differed substantially from the wide early life exploratory movements of individuals that moved to North Africa or Southern Europe.

**Appendix 3**

To assess the relationship between age and behaviour (i.e. the response variables), we modelled age using four different functional relationships: (i) a linear model, representing a consistent increase or decrease in behaviour with age; (ii) a quadratic model, characterized by a rapid initial increase in behaviour that gradually slows and stabilizes over time; (iii) an exponential model, defined by a slow initial rate of improvement that accelerates progressively over time; and (iv) a cubic model, representing an S-shaped curve with an initial slow improvement, followed by a rapid increase, and ultimately leveling off at a plateau (Hertel et al., 2023; Acácio et al., 2024).

**Table S1. GPS-tracking data used in this study.** Summary table of 101 fledglings tagged during the breeding seasons of 2015 – 2024 (last updated: 05/08/2025).

| dev | metal | PVC | device type | country | colony | orga | sex | Tagging year | included/excluded | Context |
|---|---|---|---|---|---|---|---|---|---|---|
| 6118 | CK13570 | LABZ | Interrex | France | 1 | TDV | unk | 2020 | Excluded | stop transmission |
| 8444 | CK13538 | LAVZ | Interrex | France | 1 | TDV | unk | 2021 | Included | |
| 221218 | XB330 | LBSZ | Ornitrack | France | 1 | TDV | unk | 2022 | Included | |
| 221220 | XB327 | LBST | Ornitrack | France | 1 | TDV | unk | 2022 | Included | |
| 221222 | XB576 | LCVH | Ornitrack | France | 1 | TDV | unk | 2022 | Included | |
| 221224 | XB583 | LCVV | Ornitrack | France | 1 | TDV | unk | 2022 | Included | |
| 221225 | XB329 | LBSX | Ornitrack | France | 1 | TDV | unk | 2022 | Excluded | stop transmission |
| 221226 | XB477 | LCHT | Ornitrack | France | 1 | TDV | unk | 2022 | Included | |
| 221227 | XB724 | LDLD | Ornitrack | France | 1 | TDV | unk | 2022 | Included | |
| 221228 | XB722 | LDLB | Ornitrack | France | 1 | TDV | unk | 2022 | Excluded | stop transmission |
| 221229 | XB721 | LDLA | Ornitrack | France | 1 | TDV | unk | 2022 | Included | corpse retrieved |
| 221230 | XB582 | LCVT | Ornitrack | France | 1 | TDV | Male | 2022 | Included | |
| 221231 | XB634 | LDBD | Ornitrack | France | 1 | TDV | unk | 2022 | Included | |
| 221235 | XB641 | LDBS | Ornitrack | France | 1 | TDV | unk | 2022 | Excluded | stop transmission |
| 221236 | XB468 | LCHC | Ornitrack | France | 1 | TDV | unk | 2022 | Included | |
| 221237 | XB326 | LBSS | Ornitrack | France | 1 | TDV | unk | 2022 | Included | |
| 233140 | XC405 | LJNZ | Ornitrack | France | 1 | TDV | unk | 2024 | Included | |
| 233141 | XC420 | LJPZ | Ornitrack | France | 1 | TDV | unk | 2024 | Included | |
| 233146 | XC375 | LJJZ | Ornitrack | France | 1 | TDV | unk | 2024 | Included | |
| 233147 | XC668 | LLTJ | Ornitrack | France | 1 | TDV | unk | 2024 | Included | |
| 233148 | XC435 | LJSZ | Ornitrack | France | 1 | TDV | unk | 2024 | Included | |
| 233150 | XC345 | LJFZ | Ornitrack | France | 1 | TDV | unk | 2024 | Included | |
| 233154 | XC667 | LLTL | Ornitrack | France | 1 | TDV | unk | 2024 | Included | |
| 233156 | XC465 | LJVZ | Ornitrack | France | 1 | TDV | unk | 2024 | Excluded | corpse retrieved |
| 233157 | | LHZZ | Ornitrack | France | 1 | TDV | unk | 2024 | Excluded | corpse retrieved |
| 234766 | | LJXZ | Ornitrack | France | 1 | TDV | unk | 2024 | Excluded | corpse retrieved |
| 234764 | CK13539 | LHXH | Ornitrack | France | 1 | TDV | unk | 2023 | Included | |
| 234765 | CK13590 | LHZF | Ornitrack | France | 1 | TDV | unk | 2023 | Included | |
| 234767 | CK13586 | LHZA | Ornitrack | France | 1 | TDV | unk | 2023 | Included | |

| | | | | | | | | | | |
|---|---|---|---|---|---|---|---|---|---|---|
| 234768 | CK13592 | LHZJ | Ornitrack | France | 1 | TDV | unk | 2023 | Excluded | stop transmission |
| 234769 | CK13595 | LHZP | Ornitrack | France | 1 | TDV | unk | 2023 | Included | |
| 234770 | CK13591 | LHZH | Ornitrack | France | 1 | TDV | unk | 2023 | Included | |
| 234771 | CK13599 | LHZX | Ornitrack | France | 1 | TDV | unk | 2023 | Included | |
| 234772 | CK13565 | LHXL | Ornitrack | France | 1 | TDV | unk | 2023 | Excluded | corpse retrieved |
| 234773 | CK13566 | LHXN | Ornitrack | France | 1 | TDV | unk | 2023 | Included | |
| 234772_b | | LHXZ | Ornitrack | France | 1 | TDV | unk | 2024 | Excluded | corpse retrieved |
| 234774 | XC330 | LJDZ | Ornitrack | France | 1 | TDV | unk | 2024 | Included | |
| 234775 | CK13598 | LHZV | Ornitrack | France | 1 | TDV | unk | 2023 | Included | |
| 234776 | CK13587 | LHZB | Ornitrack | France | 1 | TDV | unk | 2023 | Included | |
| 234777 | XC390 | LJLZ | Ornitrack | France | 1 | TDV | unk | 2024 | Included | |
| 234778 | CK13581 | LHXV | Ornitrack | France | 1 | TDV | unk | 2023 | Included | |
| 234779 | CK13568 | LHXS | Ornitrack | France | 1 | TDV | unk | 2023 | Included | |
| 234780 | CK13596 | LHZS | Ornitrack | France | 1 | TDV | unk | 2023 | Excluded | resident |
| 234781 | CK13593 | LHZL | Ornitrack | France | 1 | TDV | unk | 2023 | Included | |
| 234782 | CK13588 | LHZC | Ornitrack | France | 1 | TDV | unk | 2023 | Excluded | corpse retrieved |
| 234782_b | | LJHZ | Ornitrack | France | 1 | TDV | unk | 2024 | Excluded | corpse retrieved |
| 234783 | CK13567 | LHXP | Ornitrack | France | 1 | TDV | unk | 2023 | Included | |
| 234784 | CK13564 | LHXJ | Ornitrack | France | 1 | TDV | unk | 2023 | Included | |
| 234785 | CK13569 | LHXT | Ornitrack | France | 1 | TDV | unk | 2023 | Included | corpse retrieved |
| 234786 | CK13584 | LHXX | Ornitrack | France | 1 | TDV | unk | 2023 | Included | |
| 234787 | XC313 | LHZD | Ornitrack | France | 1 | TDV | unk | 2024 | Included | |
| 234788 | CK13594 | LHZN | Ornitrack | France | 1 | TDV | unk | 2023 | Included | |
| 4159A610 | E16099 | WKTV | Milsar | Italy | 2 | ISPRA | Female | 2022 | Included | |
| 76C9AC10 | E16037 | WKLS | Milsar | Italy | 2 | ISPRA | Male | 2022 | Included | |
| 78C9AC10 | E16089 | WKTD | Milsar | Italy | 2 | ISPRA | Female | 2022 | Included | |
| 7DC9AC10 | E16157 | WLBS | Milsar | Italy | 2 | ISPRA | Male | 2022 | Included | |
| 84C9AC10 | E16019 | WKKL | Milsar | Italy | 2 | ISPRA | Female | 2022 | Included | |
| FLAI01 | E14950 | WHJN | Ecotone | Italy | 2 | ISPRA | Male | 2016 | Included | |
| FLAI02 | E14954 | WHJV | Ecotone | Italy | 2 | ISPRA | Male | 2016 | Included | |
| FLAI03a | E14969 | WHKZ | Ecotone | Italy | 2 | ISPRA | Male | 2016 | Included | |
| FLAI03b | E15469 | NA | Ecotone | Italy | 2 | ISPRA | Male | 2017 | Included | |
| FLAI04 | E14980 | WHLN | Ecotone | Italy | 2 | ISPRA | Male | 2016 | Included | |
| FLAI05 | E14988 | WHNC | Ecotone | Italy | 2 | ISPRA | Male | 2016 | Included | |
| ISPR08 | E14986 | WHNA | Ecotone | Italy | 2 | ISPRA | Male | 2016 | Included | |

| | | | | | | | | | | |
|---|---|---|---|---|---|---|---|---|---|---|
| ISPR09 | E14976 | WHLH | Ecotone | Italy | 2 | ISPRA | Female | 2016 | Included | |
| ISPR10 | E14999 | WHNZ | Ecotone | Italy | 2 | ISPRA | Male | 2016 | Included | |
| ISPR11b | E15477 | NA | Ecotone | Italy | 2 | ISPRA | Male | 2017 | Included | |
| MALA11 | E14989 | WHND | Ecotone | Italy | 2 | ISPRA | Male | 2016 | Included | |
| MALA13 | E14995 | WHNP | Ecotone | Italy | 2 | ISPRA | Female | 2016 | Included | |
| MALA35 | E15495 | NA | Ecotone | Italy | 2 | ISPRA | Male | 2017 | Included | |
| SPOO02 | E14978 | WHLK | Ecotone | Italy | 2 | ISPRA | Male | 2016 | Included | |
| SPOO04 | E14998 | WHNV | Ecotone | Italy | 2 | ISPRA | Female | 2016 | Included | |
| WSIK67 | E16205 | WLFZ | Ecotone | Italy | 2 | ISPRA | Female | 2022 | Included | |
| WSIK82 | E16198 | WLFK | Ecotone | Italy | 2 | ISPRA | Male | 2022 | Included | |
| ISPR01 | E7266 | NA | Ecotone | Italy | 3 | ISPRA | Male | 2016 | Excluded | resident |
| ISPR02 | E7267 | NA | Ecotone | Italy | 3 | ISPRA | Male | 2016 | Excluded | resident |
| ISPR03 | E7268 | NA | Ecotone | Italy | 3 | ISPRA | Male | 2016 | Excluded | resident |
| ISPR04 | E7269 | NA | Ecotone | Italy | 3 | ISPRA | Female | 2016 | Included | |
| ISPR05 | E7270 | NA | Ecotone | Italy | 3 | ISPRA | Male | 2016 | Included | |
| ISPR06 | E7271 | NA | Ecotone | Italy | 3 | ISPRA | Female | 2016 | Included | |
| ISPR07 | E7272 | NA | Ecotone | Italy | 3 | ISPRA | Male | 2016 | Included | |
| MALA05 | E7281 | NA | Ecotone | Italy | 3 | ISPRA | Male | 2016 | Included | |
| MALA31 | E7265 | NA | Ecotone | Italy | 3 | ISPRA | Male | 2016 | Included | |
| SPOO01 | E7282 | NA | Ecotone | Italy | 3 | ISPRA | Male | 2016 | Included | |
| SPOO03 | E7264 | NA | Ecotone | Italy | 3 | ISPRA | Male | 2016 | Included | |
| MALA01 | E7285 | NA | Ecotone | Italy | 3 | ISPRA | Male | 2015 | Included | |
| MALA02 | E7284 | NA | Ecotone | Italy | 3 | ISPRA | Female | 2015 | Included | |
| MALA03 | E7283 | NA | Ecotone | Italy | 3 | ISPRA | Female | 2015 | Excluded | resident |
| MALA04 | E7287 | NA | Ecotone | Italy | 3 | ISPRA | Male | 2015 | Included | |
| ISPR11a | E10505 | NA | Ecotone | Italy | 4 | ISPRA | Male | 2016 | Included | |
| ISPR12 | E10506 | NA | Ecotone | Italy | 4 | ISPRA | Female | 2016 | Included | |
| ISPR13 | E10507 | NA | Ecotone | Italy | 4 | ISPRA | Male | 2016 | Included | |
| ISPR14 | E10508 | NA | Ecotone | Italy | 4 | ISPRA | Male | 2016 | Included | |
| ISPR15 | E10509 | NA | Ecotone | Italy | 4 | ISPRA | Female | 2016 | Included | |
| ISPR16 | E10510 | NA | Ecotone | Italy | 4 | ISPRA | Female | 2016 | Included | |
| ISPR17 | E10511 | NA | Ecotone | Italy | 4 | ISPRA | Male | 2016 | Included | |
| ISPR18 | E10512 | NA | Ecotone | Italy | 4 | ISPRA | Male | 2016 | Included | |
| MALA06 | E10501 | NA | Ecotone | Italy | 4 | ISPRA | Female | 2016 | Included | |
| MALA14 | E10502 | NA | Ecotone | Italy | 4 | ISPRA | Male | 2016 | Included | |

| | | | | | | | | | | |
|---|---|---|---|---|---|---|---|---|---|---|
| MALA30 | E10503 | NA | Ecotone | Italy | 4 | ISPRA | Male | 2016 | Excluded | tag failure |
| MALA34 | E10504 | NA | Ecotone | Italy | 4 | ISPRA | Male | 2016 | Included | |

*DEV = device unique identifier.
Colony numbers are shown according to Figure 1: (1) Aigues-Mortes (43.55° N, 4.18° E), (2) Molentargius (39.23° N, 9.12° E), (3) Comacchio (44.64° N, 12.21° E) and (4) Margherita di Savoia (41.39° N, 16.01° E).
Orga = Tagging organisation: Tour du Valat (TDV) and Istituto Superiore per la Protezione e la Ricerca Ambientale (ISPRA)

**Table S2.** List of traditional sites where large aggregations of greater flamingos are known to breed or spend the non-breeding season in our study area. We used Important Bird and Biodiversity Areas (IBAs) designated for greater flamingos and the most up-to-date map of flamingo breeding colony locations, developed through expert judgment during the 7th Greater Flamingo Workshop in the Mediterranean and West Africa, 2024 to map the polygons for each traditional site (Figure 3A).

| Region | Country | Name | Site ID | type |
|---|---|---|---|---|
| Africa | Algeria | Chott Merouane et Oued Khrouf | 6174 | flamingo.iba |
| Africa | Algeria | Garaet et-Tarf | 6169 | flamingo.iba |
| Africa | Algeria | Sebkha d'Oran | 6173 | flamingo.iba |
| Africa | Algeria | Sebkhet Baker | 6165 | flamingo.iba |
| Africa | Algeria | Sebkhet Djendli | 6168 | flamingo.iba |
| Africa | Egypt | El Malaha | 6189 | flamingo.iba |
| Africa | Egypt | Lake Bardawil | 6187 | flamingo.iba |
| Africa | Egypt | Zaranik Protected Area | 6188 | flamingo.iba |
| Africa | Libya | Benghazi | 6472 | flamingo.iba |
| Africa | Mauritania | Aftout es Sâheli | 6634 | flamingo.iba |
| Africa | Mauritania | Banc d'Arguin National Park | 6629 | flamingo.iba |
| Africa | Mauritania | Diawling National Park | 6643 | flamingo.iba |
| Africa | Morocco | Baie d'Ad Dakhla | 6520 | flamingo.iba |
| Africa | Morocco | Lagune de Khnifiss | 6517 | flamingo.iba |
| Africa | Morocco | Marais Larache | 6482 | flamingo.iba |
| Africa | Morocco | Merja Zerga | 6487 | flamingo.iba |
| Africa | Morocco | Sebkha Bou Areg | 6483 | flamingo.iba |
| Africa | Morocco | Sebkha Zima | 6505 | flamingo.iba |
| Africa | Tunisia | Chott Djerid | 6952 | flamingo.iba |

| | | | | |
|---|---|---|---|---|
| Africa | Tunisia | Côte de Zerkine et El Grine | 6954 | flamingo.iba |
| Africa | Tunisia | Côtes de l'Île de Djerba | 6953 | flamingo.iba |
| Africa | Tunisia | Garaet Douza | 6946 | flamingo.iba |
| Africa | Tunisia | Golfe de Boughrara | 6955 | flamingo.iba |
| Africa | Tunisia | Ichkeul | 6919 | flamingo.iba |
| Africa | Tunisia | Lac de Tunis | 6926 | flamingo.iba |
| Africa | Tunisia | Lagune El Bibane | 6961 | flamingo.iba |
| Africa | Tunisia | Salines de Monastir | 6938 | flamingo.iba |
| Africa | Tunisia | Salines de Thyna | 6944 | flamingo.iba |
| Africa | Tunisia | Sebkhet Ennoual | 6948 | flamingo.iba |
| Africa | Tunisia | Sebkhet Halk el Menzel | 6934 | flamingo.iba |
| Africa | Tunisia | Sebkhet Kelbia | 6937 | flamingo.iba |
| Africa | Tunisia | Sebkhet Sejoumi | 6927 | flamingo.iba |
| Africa | Tunisia | Sebkhet Sidi El Hani | 6941 | flamingo.iba |
| Africa | Tunisia | Sebkhet Sidi Khelifa | 6933 | flamingo.iba |
| Africa | Tunisia | Sebkhet Sidi Mansour | 6950 | flamingo.iba |
| Africa | Tunisia | Îles Kerkennah | 6943 | flamingo.iba |
| Africa | Tunisia | Îles Kneiss | 6949 | flamingo.iba |
| Europe | Cyprus | Akrotiri Peninsula - Episkopi Cliffs | 24489 | flamingo.iba |
| Europe | Cyprus | Larnaca Salt Lakes | 30 | flamingo.iba |
| Europe | France | Camargue | 2327 | flamingo.iba |
| Europe | France | Cordon lagunaire de Sète à Agde | 2298 | flamingo.iba |
| Europe | France | Etangs Montpellierains | 2291 | flamingo.iba |
| Europe | France | Etangs Narbonnais | 2286 | flamingo.iba |
| Europe | France | Etangs de Canet et de Villeneuve-de-la-Raho et embouchure du Tech | 2302 | flamingo.iba |
| Europe | France | Etangs de Citis, Lavalduc, Engrenier, Pourra, l'Estomac, Fos et salines de Rassuen et de Fos | 2340 | flamingo.iba |
| Europe | France | Etangs de Leucate et Lapalme | 2285 | flamingo.iba |
| Europe | France | Petite Camargue fluvio-lacustre | 2305 | flamingo.iba |
| Europe | France | Petite Camargue laguno-marine | 2306 | flamingo.iba |
| Europe | France | Salins d'Hyères et de Pesquiers | 2335 | flamingo.iba |
| Europe | Greece | Evros delta | 986 | flamingo.iba |
| Europe | Greece | Kalloni Gulf, Lesvos | 1086 | flamingo.iba |
| Europe | Greece | Lake Ismarida (Mitrikou) | 988 | flamingo.iba |

| | | | | |
|---|---|---|---|---|
| Europe | Greece | Lakes Chortarolimni and Alyki, Moudros gulf, Diapori fen and Fakos peninsula, Limnos | 1085 | flamingo.iba |
| Europe | Greece | Lakes Volvi, Koroneia and Rentina Gorge | 1006 | flamingo.iba |
| Europe | Greece | Porto Lagos, lake Vistonida and coastal lagoons | 989 | flamingo.iba |
| Europe | Italy | Argentario, Laguna di Orbetello Lagoon e Lago di Burano | 9582 | flamingo.iba |
| Europe | Italy | Cagliari wetlands | 2824 | flamingo.iba |
| Europe | Italy | Colostrai pool | 2822 | flamingo.iba |
| Europe | Italy | Gargano Promontory and Capitanata Wetlands | 9583 | flamingo.iba |
| Europe | Italy | Palmas Gulf wetlands | 2846 | flamingo.iba |
| Europe | Italy | Sinis and Oristano wetlands | 9585 | flamingo.iba |
| Europe | Italy | Tarquinia saltpans | 2882 | flamingo.iba |
| Europe | Montenegro | Ulcinj saltpans | 117 | flamingo.iba |
| Europe | Portugal | Castro Marim | 378 | flamingo.iba |
| Europe | Portugal | Mondego Estuary | 19672 | flamingo.iba |
| Europe | Portugal | Ria Formosa (Faro lagoon) | 377 | flamingo.iba |
| Europe | Portugal | Salinas de Alverca e do Forte da Casa | 19678 | flamingo.iba |
| Europe | Portugal | Tejo estuary | 370 | flamingo.iba |
| Europe | Spain | Cádiz bay | 1859 | flamingo.iba |
| Europe | Spain | Ebro delta | 1811 | flamingo.iba |
| Europe | Spain | El Hondo wetland | 1824 | flamingo.iba |
| Europe | Spain | Fuente de Piedra, Gosque and Campillos lakes | 1875 | flamingo.iba |
| Europe | Spain | Guadalquivir marshes | 1855 | flamingo.iba |
| Europe | Spain | Medina and Puerto Real lagoons | 1860 | flamingo.iba |
| Europe | Spain | Mountain range and saltpans at Cabo de Gata | 1835 | flamingo.iba |
| Europe | Spain | Odiel and Tinto marshes and Huelva coastal lagoons | 1854 | flamingo.iba |
| Europe | Spain | Santa Pola salt-pans | 1825 | flamingo.iba |
| Europe | Spain | Wetlands of western Almería | 1838 | flamingo.iba |
| Europe | Türkiye | Acigöl Lake | 763 | flamingo.iba |
| Europe | Türkiye | Burdur Lake | 766 | flamingo.iba |
| Europe | Türkiye | Büyük Menderes Delta | 760 | flamingo.iba |
| Europe | Türkiye | Ceyhan Delta | 24063 | flamingo.iba |
| Europe | Türkiye | Eregli Plain | 749 | flamingo.iba |
| Europe | Türkiye | Gediz Delta | 787 | flamingo.iba |
| Europe | Türkiye | Göksu Delta | 768 | flamingo.iba |
| Europe | Türkiye | Gökçeada Lagoon | 9628 | flamingo.iba |

| | | | | |
|---|---|---|---|---|
| Europe | Türkiye | Kulu Lake | 752 | flamingo.iba |
| Europe | Türkiye | Seyfe Lake | 754 | flamingo.iba |
| Europe | Türkiye | Seyhan Delta | 31658 | flamingo.iba |
| Europe | Türkiye | Sultan Marsh | 755 | flamingo.iba |
| Europe | Türkiye | Tersakan Lake | 791 | flamingo.iba |
| Europe | Türkiye | Tuz Lake | 757 | flamingo.iba |
| Europe | Türkiye | Çöl Lake and Çalikdüzü | 807 | flamingo.iba |
| Europe | Italy | Valli di Comacchio e Bonifica del Mezzano | 2750 | breeding colony |
| Europe | Spain | S'Albufera de Mallorca y S'Albufereta de Pollença | 1887 | breeding colony |
| Europe | Spain | Gallocanta | 1822 | breeding colony |
| Europe | Italy | Laguna di Venezia | 2739 | breeding colony |
| Europe | Italy | Pantani di Vendicari e di Capo Passero | 2838 | breeding colony |
| Europe | Türkiye | Akşehir ve Eber Gölleri | 747 | breeding colony |
| Africa | Algeria | Sebkhet Ez-Zemoul | 6167 | breeding colony |
| Europe | Italy | Diaccia Botrona | 2765 | breeding colony |

**Table S3:** Summary of the number of flamingos tagged as fledglings in Aigues-Mortes (France) and Molentargius, Comacchio and Margherita di Savoia (Italy). Individuals of unknown sex across all age classes are also shown. To assess the effect of sex on residency metrics, unknown sex and older individuals (i.e., 5 years and older) were excluded due to the tiny sample size.

| Age | sex | Number of individuals |
|---|---|---|
| 1yo | Female | 10 |
| 1yo | Male | 35 |
| 1yo | unk | 37 |
| 2yo | Female | 8 |
| 2yo | Male | 20 |
| 2yo | unk | 24 |
| 3yo | Female | 6 |
| 3yo | Male | 17 |
| 3yo | unk | 8 |
| 4yo | Female | 5 |
| 4yo | Male | 14 |
| 5yo | Female | 3 |
| 5yo | Male | 10 |
| 6yo | Female | 2 |
| 6yo | Male | 7 |
| 7yo | Female | 1 |
| 7yo | Male | 4 |
| 8yo | Male | 2 |
| 8yo | Male | 0 |

**Table S4.** Results of the performance of GLMMs comparing the interaction effect of season (with two levels: autumn-winter, summer-spring) and age, modelled as a linear, quadratic, exponential, or polynomial effect, used to explain the variation in the number of staging events, staging duration (days), distance of main sites to the natal colony, distance from previous main sites (km) and the distance between consecutive staging events. The performance of the interaction effect of traditional sites (with two levels: inside and outside) and the different functional relationships of age used to explain the variation in segment duration is also shown. The proportion of variance explained by fixed ($R^2_{marginal}$) and random ($R^2_{conditional}$) factors are given. The model with the lowest AIC was preferred. When the top models had a ΔAIC ≤ 2 units, we retained the simplest model. Models selected based on these criteria are highlighted in bold.

| | AIC | delta AIC | Weight | $R^2_{Conditional}$ | $R^2_{Marginal}$ |
|---|---|---|---|---|---|
| **Number of staging events** | | | | | |
| **age lineal** | **2265.31** | **0.00** | **0.51** | **0.44** | **0.13** |
| age quadratic | 2267.23 | 1.91 | 0.20 | 0.45 | 0.13 |
| age exponential | 2267.47 | 2.16 | 0.17 | 0.47 | 0.12 |
| age cubic | 2268.24 | 2.93 | 0.12 | 0.44 | 0.12 |
| **Average staging duration (days)** | | | | | |
| age exponential | 3561.67 | 0.00 | 0.56 | 0.34 | 0.08 |
| **age lineal** | **3563.16** | **1.49** | **0.27** | **0.32** | **0.10** |
| age cubic | 3564.62 | 2.96 | 0.13 | 0.33 | 0.10 |
| age quadratic | 3566.78 | 5.11 | 0.04 | 0.33 | 0.10 |
| **Distance of main sites to the natal colony** | | | | | |
| **age lineal** | **3189.34** | **0.00** | **0.52** | **0.75** | **0.03** |
| age cubic | 3191.15 | 1.81 | 0.21 | 0.76 | 0.04 |
| age quadratic | 3191.70 | 2.36 | 0.16 | 0.75 | 0.03 |
| age exponential | 3192.28 | 2.94 | 0.12 | 0.75 | 0.02 |
| **Distance from previous main sites (km)** | | | | | |
| age cubic | 2302.59 | 0.00 | 0.55 | 0.43 | 0.07 |
| **age lineal** | **2303.79** | **1.20** | **0.30** | **0.42** | **0.04** |
| age exponential | 2306.44 | 3.86 | 0.08 | 0.41 | 0.04 |
| age quadratic | 2306.78 | 4.19 | 0.07 | 0.41 | 0.04 |
| **Average segment duration (days)** | | | | | |
| **age lineal** | **6281.96** | **0.00** | **0.68** | **0.25** | **0.13** |
| age quadratic | 6285.09 | 3.13 | 0.14 | 0.26 | 0.12 |
| age exponential | 6285.11 | 3.15 | 0.14 | 0.26 | 0.13 |
| age polynomial | 6288.06 | 6.10 | 0.03 | 0.26 | 0.13 |

**Table S5. Summary of the global GLMMs testing the effect of season and age on residency metrics of greater flamingos during their early life.** Model estimates (SE) and 95% confidence interval for the parameters retained in the best model, random effects (ID, year and colony), the proportion of variance explained by fixed ($R^2_{marginal}$) and random ($R^2_{conditional}$) factors and sample size, in parenthesis, are given. Parameter estimates with confidence intervals that do not overlap zero are shown in bold.

| | | | Fixed effects | | | Random effects | | |
|---|---|---|---|---|---|---|---|---|
| | **$R^2_{mar}$** | **$R^2_{con}$** | | **Estimate (SE)** | **95% CI** | | **Var** | **SD** |
| Number of staging events (n=377, individuals = 83) | 0.15 | 0.60 | **Intercept** | **1.48 (0.07)** | **1.35, 1.61** | Individual | 0.21 | 0.46 |
| | | | **age linear** | **-0.23 (0.04)** | **-0.31, -0.15** | | | |
| | | | **season** | **-0.16 (0.05)** | **-0.25, -0.06** | | | |
| | | | age linear x season | -0.05 (0.05) | -0.16, 0.06 | | | |
| Average staging duration (days) (n=377, individuals = 83) | 0.08 | 0.34 | **Intercept** | **41.66 (6.58)** | **28.72, 54.59** | Individual | 287.73 | 16.96 |
| | | | **age lineal** | **7.99 (3.49)** | **1.12, 14.85** | Year | 113.16 | 10.64 |
| | | | **season** | **17.12 (4.05)** | **9.13, 25.11** | Colony | 74.67 | 8.64 |
| | | | age linear x season | 4.14 (4.05) | -3.83, 12.11 | Residual | 1394.73 | 37.35 |
| Distance of principal sites to the natal colony (km) (n=231, individuals =53) | 0.03 | 0.75 | **Intercept** | 6 (0.17) | 5.67, 6.33 | Individual | 1.18 | 1.08 |
| | | | age lineal | -0.14 (0.07) | -0.28, -0.01 | | | |
| | | | **season** | **-0.2 (0.09)** | **-0.38, -0.03**[ms] | | | |
| | | | **age linear x season** | -0.1 (0.09) | -0.28, 0.08 | | | |
| Distance from previous principal sites (km) (n=231, individuals =53) | 0.09 | 0.31 | **Intercept** | **4.20 (0.29)** | **3.64, 4.77** | Individual | 1.77 | 1.33 |
| | | | age lineal | -0.23 (0.19) | -0.61, 0.15 | | | |
| | | | **season** | **0.53 (0.28)** | **-0.01, 1.07**[ms] | | | |
| | | | **age linear x season** | **0.67 (0.26)** | **0.15, 1.18** | | | |
| Mean segment duration (days) (n=960, individuals = 83) | 0.03 | 0.25 | Intercept | 28.33 (6.43) | 15.72, 40.93 | Individual | 354.74 | 18.83 |
| | | | **age exponential** | **2.84 (0.68)** | **1.51, 4.17** | Year | 45.24 | 6.72 |
| | | | TS | 1.24 (2.81) | -4.27, 6.75 | Colony | 99.80 | 9.99 |
| | | | season | 3.58 (2.81) | -1.93, 9.09 | Residual | 1657.02 | 40.70 |
| | | | **age exponential x TS** | **-3.68 (0.88)** | **-5.41, -1.95** | | | |

*For the categorical variable, parameter estimates are given relative to the reference category (season: spring-summer and traditional sites: outside).

The effect of age was explored as linear, quadratic, exponential and cubic functional relationships. For all the residency metrics the linear function was preferred (see Table S2).

ms= marginal significant

TS = Traditional sites

**Table S6.** Details of top-ranked models explaining variation in the number of staging events, average staging duration (days), distance of main sites to the natal colony (km) and distance from previous main sites (km), including age as a linear effect and their interaction with season as explanatory variables. Models are ranked according to increasing AIC values, with the best-performing model on top. Grey boxes indicate that a given variable is included in the model. Age is scaled and centred.

| **Response** | **Intercept** | **Season** | **Age** | **Interaction** | **df** | **AIC** | **delta** | **weight** | **$R^2_{Mar}$** | **$R^2_{Con}$** |
|---|---|---|---|---|---|---|---|---|---|---|
| Number of staging events | | | | | | | | | | |
| | 1.473732 | | | | 4 | 1830.738 | 0 | 0.615 | 0.15 | 0.59 |
| | 1.481209 | | | | 5 | 1831.767 | 1.02 | 0.368 | 0.15 | 0.59 |
| | 1.42744 | | | | 3 | 1838.022 | 7.28 | 0.016 | 0.13 | 0.58 |
| | 1.571732 | | | | 3 | 1894.04 | 63.30 | 1.11E-14 | 0.001 | 0.52 |
| | 1.529213 | | | | 2 | 1898.546 | 67.80 | 1.16E-15 | 0 | 0.51 |
| Average staging duration (days) | 41.65854 | | | | 8 | 3857.1 | 0 | 0.862 | 0.09 | 0.32 |
| | 41.70056 | | | | 7 | 3860.7 | 3.67 | 0.138 | 0.09 | 0.32 |
| | 42.82629 | | | | 6 | 3871.7 | 14.6 | 6E-04 | 0.02 | 0.36 |
| | 49.02043 | | | | 6 | 3880.6 | 23.5 | 7E-06 | 0.03 | 0.31 |
| | 48.92126 | | | | 5 | 3887.1 | 30.1 | 3E-07 | 0 | 0.38 |
| Distance of main sites to natal the colony (km) | 5.991811 | | | | 5 | 3188.6 | 0 | 0.531 | 0.03 | 0.75 |
| | 6.000641 | | | | 6 | 3189.3 | 0.74 | 0.367 | 0.03 | 0.75 |
| | 5.933366 | | | | 4 | 3192 | 3.4 | 0.097 | 0.02 | 0.74 |
| | 6.072138 | | | | 4 | 3198.6 | 10 | 0.004 | 0.01 | 0.73 |
| | 6.016208 | | | | 3 | 3200.6 | 12 | 0.001 | 0 | 0.73 |
| Distance from previous main sites (km) | 4.203434 | | | | 6 | 2303.8 | 0 | 0.626 | 0.04 | 0.42 |
| | 4.279859 | | | | 4 | 2306.4 | 2.63 | 0.168 | 0.01 | 0.4 |
| | 4.504295 | | | | 3 | 2307.7 | 3.88 | 0.09 | 0 | 0.36 |
| | 4.300072 | | | | 5 | 2308.1 | 4.3 | 0.073 | 0.01 | 0.41 |
| | 4.519883 | | | | 4 | 2309.2 | 5.37 | 0.043 | 0 | 0.38 |

**Table S7. Summary of the final GLMMs testing the effect of season and age on staging metrics of greater flamingos during their early life.** Model estimates (SE) and 95% confidence interval for the parameters retained in the best model, random effects (ID, year and colony), the proportion of variance explained by fixed ($R^2_{marginal}$) and random ($R^2_{conditional}$) factors and sample size, in parenthesis, are given. Parameter estimates with confidence intervals that do not overlap zero are shown in bold.

| | | | Fixed effects | | | | Random effects | |
|---|---|---|---|---|---|---|---|---|
| | $R^2_{mar}$ | $R^2_{con}$ | | Estimate (SE) | 95% CI | | Variance | SD |
| Number of staging events | 0.15 | 0.60 | **Intercept** | **1.47 (0.07)** | **1.34, 1.60** | Individual | 0.21 | 0.46 |
| (n=377, individuals = 83) | | | **age linear** | **-0.25 (0.03)** | **-0.32, -0.19** | | | |
| | | | **season** | **-0.14 (0.05)** | **-0.24, -0.05** | | | |
| Average stage duration in staging sites (days) | 0.08 | 0.34 | **Intercept** | **41.66 (6.58)** | **28.72, 54.59** | Individual | 287.73 | 16.96 |
| (n=377, individuals = 83) | | | **age lineal** | **7.99 (3.49)** | **1.12, 14.85** | Year | 113.16 | 10.64 |
| | | | **season** | **17.12 (4.05)** | **9.13, 25.11** | Colony | 74.67 | 8.641 |
| | | | age linear x season | 4.14 (4.05) | -3.83, 12.11 | Residual | 1394.73 | 37.35 |
| Distance of main sites to natal the colony (km) | 0.03 | 0.75 | **Intercept** | **5.99 (0.17)** | **5.67, 6.32** | Individual | 1.17 | 1.08 |
| (n=231, individuals =53) | | | **age lineal** | **-0.19 (0.05)** | **-0.30, -0.08** | | | |
| | | | **season** | **-0.21 (0.09)** | **-0.39, -0.03** | | | |
| Distance from previous main sites (km) | 0.09 | 0.31 | **Intercept** | **4.20 (0.29)** | **3.64, 4.77** | Individual | 1.78 | 1.33 |
| (n=231, individuals =53) | | | age lineal | -0.23 (0.19) | -0.61, 0.15 | | | |
| | | | **season** | **0.53 (0.28)** | **-0.01, 1.07**[ms] | | | |
| | | | **age linear x season** | **0.67 (0.26)** | **0.15, 1.18** | | | |
| Mean segment duration | 0.03 | 0.26 | **Intercept** | **29.58 (6.52)** | **16.79, 42.36** | Individual | 352.72 | 18.78 |
| (n=960, individuals = 83) | | | **age exponential** | **2.80 (0.68)** | **1.47, 4.13** | Year | 47.92 | 6.92 |
| | | | TS | 1.17 (2.81) | -4.35, 6.68 | Colony | 107.44 | 10.36 |
| | | | **age exponential x TS** | **-3.69 (0.88)** | **-5.41, -1.96** | Residual | 1659.51 | 40.73 |

*For the categorical variable, parameter estimates are given relative to the reference category (season: spring-summer and traditional sites: outside).
TS = traditional sites
The relationship between age and the staging metrics was analyzed using linear, quadratic, exponential, and cubic functions. Among these, the linear function was preferred for all metrics, except for mean segment duration (see Table S2)
ms= marginal significant

**Table S8.** Results of the performance of GLMMs comparing the interaction effect of sex (with two levels: males and females) and age, modelled as a linear, quadratic, exponential, or polynomial effect, used to explain the variation in the number of staging events, staging duration (days), distance of main sites to the natal colony and distance from previous main sites (km). The performance of the interaction effect of traditional sites (with two levels: inside and outside) and the different functional relationships of age used to explain the variation in segment duration is also shown. The proportion of variance explained by fixed ($R^2_{marginal}$) and random ($R^2_{conditional}$) factors are given. The model with the lowest AIC was preferred. When the top models had a ΔAIC ≤ 2 units, we retained the simplest model (i.e., the one with the fewest parameters). Models selected based on these criteria are highlighted in bold.

| | Response | | | | |
|---|---|---|---|---|---|
| | **Number of staging events** | | | | |
| | AIC | delta AIC | Weight | $R^2_{Conditional}$ | $R^2_{Marginal}$ |
| **age quadratic** | **1294.34** | **0.00** | **0.83** | **0.50** | **0.13** |
| age cubic | 1298.05 | 3.72 | 0.13 | 0.49 | 0.14 |
| age lineal | 1300.43 | 6.09 | 0.04 | 0.49 | 0.11 |
| age exponential | 1305.97 | 11.63 | 0.00 | 0.48 | 0.07 |
| | **Average staging duration (days)** | | | | |
| **age quadratic** | **2197.94** | **0.00** | **0.97** | **0.39** | **0.18** |
| age cubic | 2205.06 | 7.12 | 0.03 | 0.36 | 0.13 |
| age exponential | 2208.92 | 10.98 | 0.00 | 0.36 | 0.08 |
| age lineal | 2209.83 | 11.90 | 0.00 | 0.32 | 0.10 |
| | **Distance of main sites to the natal colony** | | | | |
| **age lineal** | **1790.138** | **0.00** | **0.443** | **0.849** | **0.017** |
| age exponential | 1790.19 | 0.05 | 0.432 | 0.849 | 0.017 |
| age quadratic | 1794.06 | 3.93 | 0.062 | 0.850 | 0.017 |
| age cubic | 1794.06 | 3.93 | 0.062 | 0.850 | 0.017 |
| | **Distance from previous main sites (km)** | | | | |
| **age lineal** | **1341.48** | **0.00** | **0.549** | **0.44** | **0.09** |
| age exponential | 1343.22 | 1.74 | 0.23 | 0.43 | 0.06 |
| age quadratic | 1344.69 | 3.21 | 0.11 | 0.41 | 0.08 |
| age cubic | 1344.69 | 3.21 | 0.11 | 0.41 | 0.08 |
| | **Average segment duration (days)** | | | | |
| **age exponential** | **5766.58** | **0.00** | **0.63** | **0.32** | **0.02** |
| age lineal | 5768.68 | 3.02 | 0.22 | 0.32 | 0.01 |
| age quadratic | 5769.70 | 3.98 | 0.13 | 0.30 | 0.02 |
| age cubic | 5773.34 | 7.52 | 0.02 | 0.30 | 0.02 |

**Table S9. Summary of the global GLMMs testing the effect of season, sex and age on staging metrics of greater flamingos during their early life.** Model estimates (SE) and 95% confidence interval for the parameters retained in the best model, random effects (ID, year and colony), the proportion of variance explained by fixed ($R^2_{marginal}$) and random ($R^2_{conditional}$) factors and sample size, in parenthesis, are given. Parameter estimates with confidence intervals that do not overlap zero are shown in bold.

| | | | Fixed effects | | | Random effects | | |
|---|---|---|---|---|---|---|---|---|
| | **$R^2_{mar}$** | **$R^2_{con}$** | | **Estimate (SE)** | **95% CI** | | **Variance** | **SD** |
| Number of staging events (n = 214, individuals = 45) | 0.18 | 0.50 | **Intercept** | **0.96 (0.19)** | **0.60, 1.33** | Individual | 0.15 | 0.38 |
| | | | **sex** | **0.54 (0.21)** | **0.13, 0.95** | | | |
| | | | **age 1st** | **-0.38 (0.08)** | **-0.55, -0.22** | | | |
| | | | **age 2nd** | **0.24 (0.10)** | **0.10, 0.48** | | | |
| | | | **season** | **-0.31 (0.07)** | **-0.44, -0.18** | | | |
| | | | **age 1st x sex** | **0.25 (0.09)** | **0.06, 0.42** | | | |
| | | | **age 2nd x sex** | **-0.30 (0.11)** | **-0.52, -0.09** | | | |
| Average staging duration (days) (n = 214, individuals = 45) | 0.119 | 0.376 | **Intercept** | **59.73 (12.22)** | **35.65, 83.82** | Individual | 355.1 | 18.84 |
| | | | $sex_{(male)}$ | -19.51 (11.89) | -42.96, 3.94 | Year | 195.6 | 13.98 |
| | | | age 1st | 10.87 (7.47) | -3.86, 25.29 | Residual | 1341.6 | 36.63 |
| | | | **age 2nd** | **-20.75 (6.97)** | **-34.49, -7.02** | | | |
| | | | **season** | **26.65 (5.78)** | **15.26, 38.04** | | | |
| | | | age 1st x sex | -8.36 (7.14) | -22.43, 5.71 | | | |
| | | | **age 2nd x sex** | **18.87 (7.59)** | **3.90, 33.83** | | | |
| Distance of principal sites to the natal colony (km) (n = 131, individuals = 28) | 0.017 | 0.849 | **Intercept** | 6.23 (0.36) | 5.52, 6.94 | Individual | 0.946 | 0.972 |
| | | | sex | -0.22 (0.43) | -1.05, 0.62 | | | |
| | | | age lineal | -0.14 (0.15) | -0.43, 0.16 | | | |
| | | | **season** | -0.06 (0.07) | -0.20, 0.09 | | | |
| | | | age linear x season | 0.2 (0.17) | -0.14, 0.54 | | | |
| Distance from previous principal sites (km) (n = 131, individuals = 28) | 0.085 | 0.439 | **Intercept** | **3.74 (0.64)** | **2.47, 5.00** | Individual | 1.233 | 1.11 |
| | | | sex | 0.62 (0.63) | -0.61, 1.85 | Year | 0.293 | 0.541 |
| | | | **age lineal** | **-0.81 (0.43)** | **-1.66, 0.04**[ms] | | | |
| | | | season | 0.45 (0.37) | -0.29, 1.18 | | | |
| | | | age x sex | 0.48 (0.40) | -0.30, 1.26 | Individual | 209.10 | 14.46 |
| Mean segment duration (days) (n=548, individuals 45) | 0.02 | 0.37 | **Intercept** | 48.39 (12.97) | 22.92, 73.86 | Year | 821.48 | 28.66 |
| | | | sex | -2.85 (7.24) | -17.07, 11.38 | Colony | 67.07 | 8.19 |
| | | | **age exponential** | **-6.23 (3.09)** | **-12.30, -0.16** | Residual | 1897.60 | 43.56 |
| | | | TS | -4.40 (3.83) | -11.93, 3.13 | | | |
| | | | season | 2.61 (4.03) | -5.31, 10.53 | | | |
| | | | age exponential x sex | 3.47 (2.62) | -1.67, 8.61 | | | |

*For the categorical variable, parameter estimates are given relative to the reference category (season: spring-summer and TS: outside, sex: male). Because we did not have enough sample size for the older age classes, we just included age 1 to age 4. Unknown sex was excluded. The effect of age was explored as linear, quadratic, exponential and cubic functional relationships (see Table S7).
ms= marginal significant

**Table S10**. Details of top-ranked models explaining variation in segment duration, including age as an exponential effect, season, traditional sites (TS = two-level categorical variables: inside, outside) and their interaction with age as explanatory variables. Models are ranked according to increasing AIC values, with the best-performing model on top. Grey boxes indicate that a given variable is included in the model. Age is scaled and centred.

| | **Intercept** | **TS** | **Season** | **Age** | **Interaction** | **df** | **AIC** | **delta** | **weight** | **$R^2_{Mar}$** | **$R^2_{Con}$** |
|---|---|---|---|---|---|---|---|---|---|---|---|
| Mean segment duration | 29.577543 | | | | | 8 | 9962.48 | 0.00 | 0.55 | 0.02 | 0.25 |
| | 28.325285 | | | | | 9 | 9962.86 | 0.38 | 0.45 | 0.02 | 0.25 |
| | 30.093706 | | | | | 6 | 9976.40 | 13.92 | 0.00 | 0.01 | 0.24 |
| | 28.846467 | | | | | 7 | 9976.71 | 14.23 | 0.00 | 0.01 | 0.24 |
| | 31.030531 | | | | | 7 | 9977.84 | 15.36 | 0.00 | 0.01 | 0.24 |

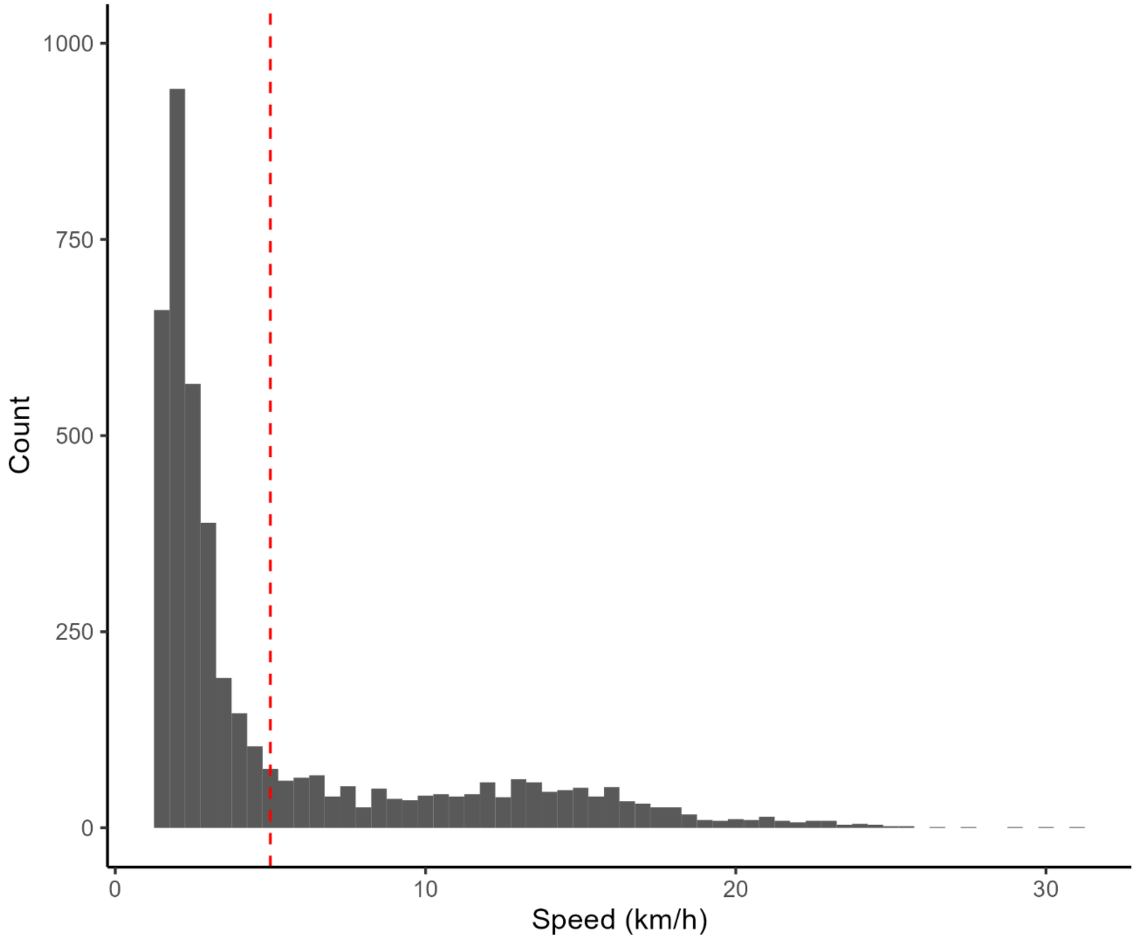


**Figure S1.** Histogram of ground speed to classify flight vs staging movements. Red vertical dashed line shows the selected cut-off used to differentiate periods of resting (< 5 km/h) and travel (>= 5 km/h).

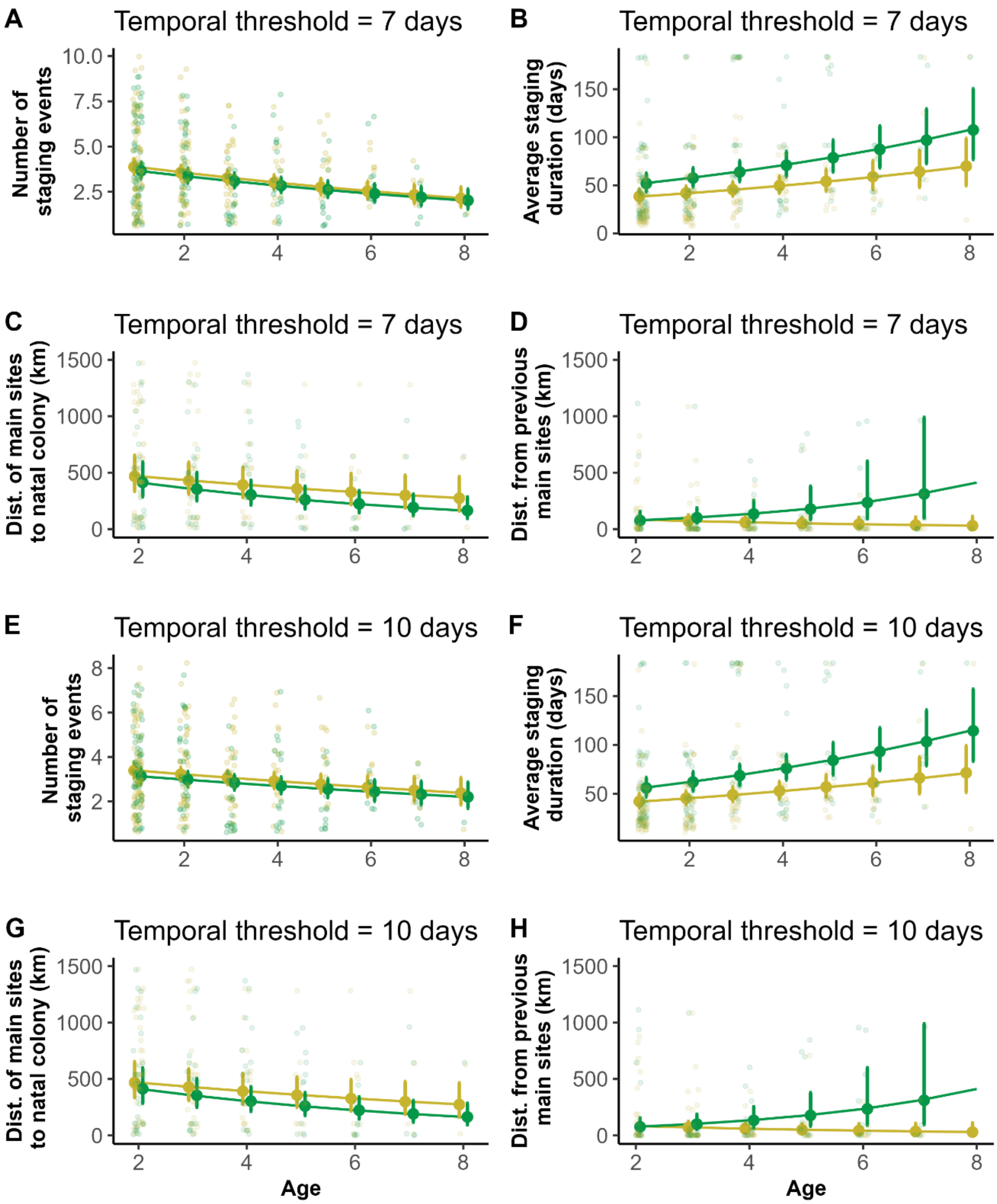


**Figure S2.** Sensitivity analyses define long residency events. In the main analyses, we use 3-day threshold. We re-ran the entire analysis using 7 days (A, B, C, D) and 10 days (E, F, G, H) and found no difference in the results.

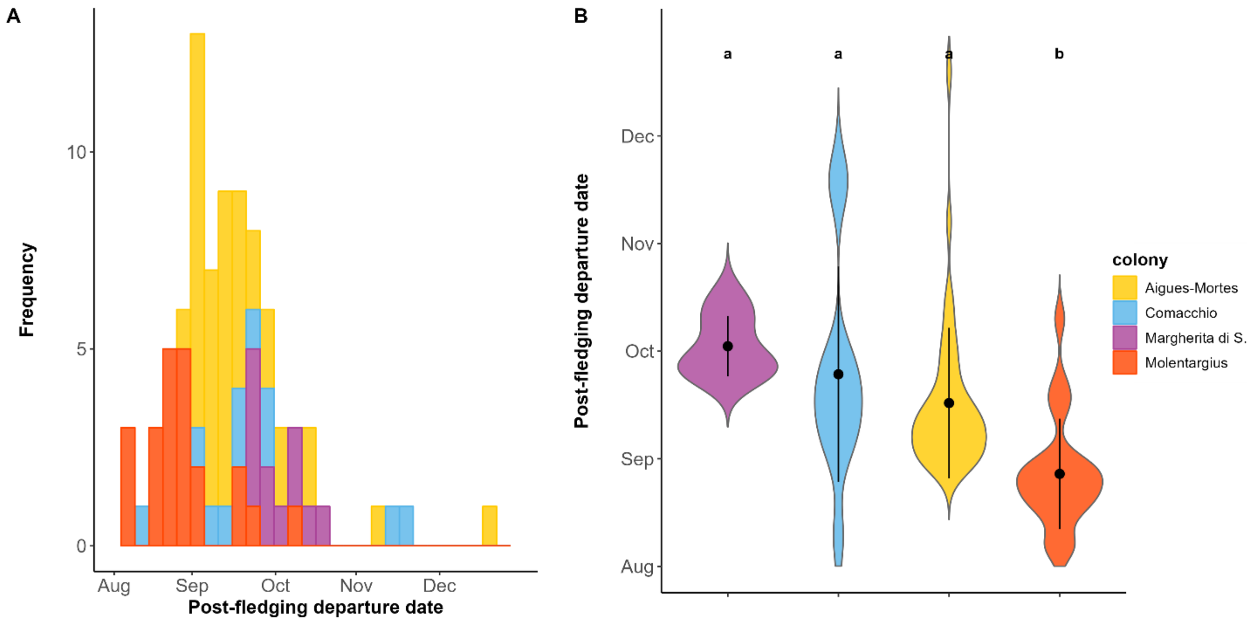


**Figure S3.** (A) Histograms of the post fledging departure date of Greater flamingos from different natal colonies: Aigues-Mortes in yellow, Molentargius in orange red, Comacchio in blue and Margherita di Savoia in purple.  (B) Violin plots illustrating the differences between post-fledging departure date by different colonies. The letters above represent significant differences by Tukey HSD post-hoc tests at the 0.05 significance level. Black dots and whiskers indicate the mean ± sd in each colony. Groups sharing the same letter are not significantly different.

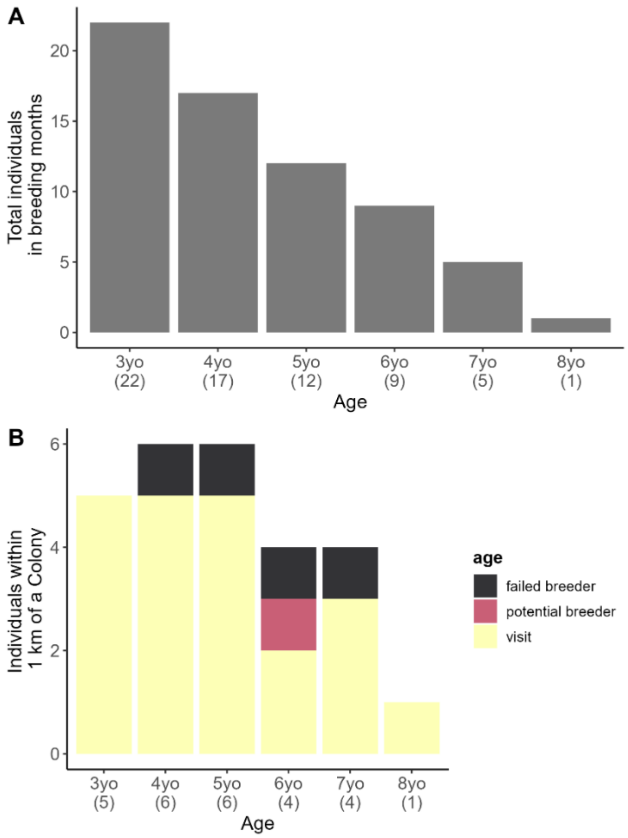


**Figure S4. Age distribution of birds displaying possible breeding behaviour.** (A) Physiologically mature individuals (3yo) tracked during the breeding months (April, May, June) (N = 66 year-age combinations). The number of tracked birds decreased over time due to death, loss, or failure of the GPS transmitters. (B) A subset showing the number of individuals within a 1 km radius of a known flamingo colony, displaying possible breeding behaviour. The colours represent individuals classified as "failed breeders" (black), "potential breeders" (fuchsia), and "visits" (yellow). The numbers of unique individuals tracked each year are indicated in parentheses.

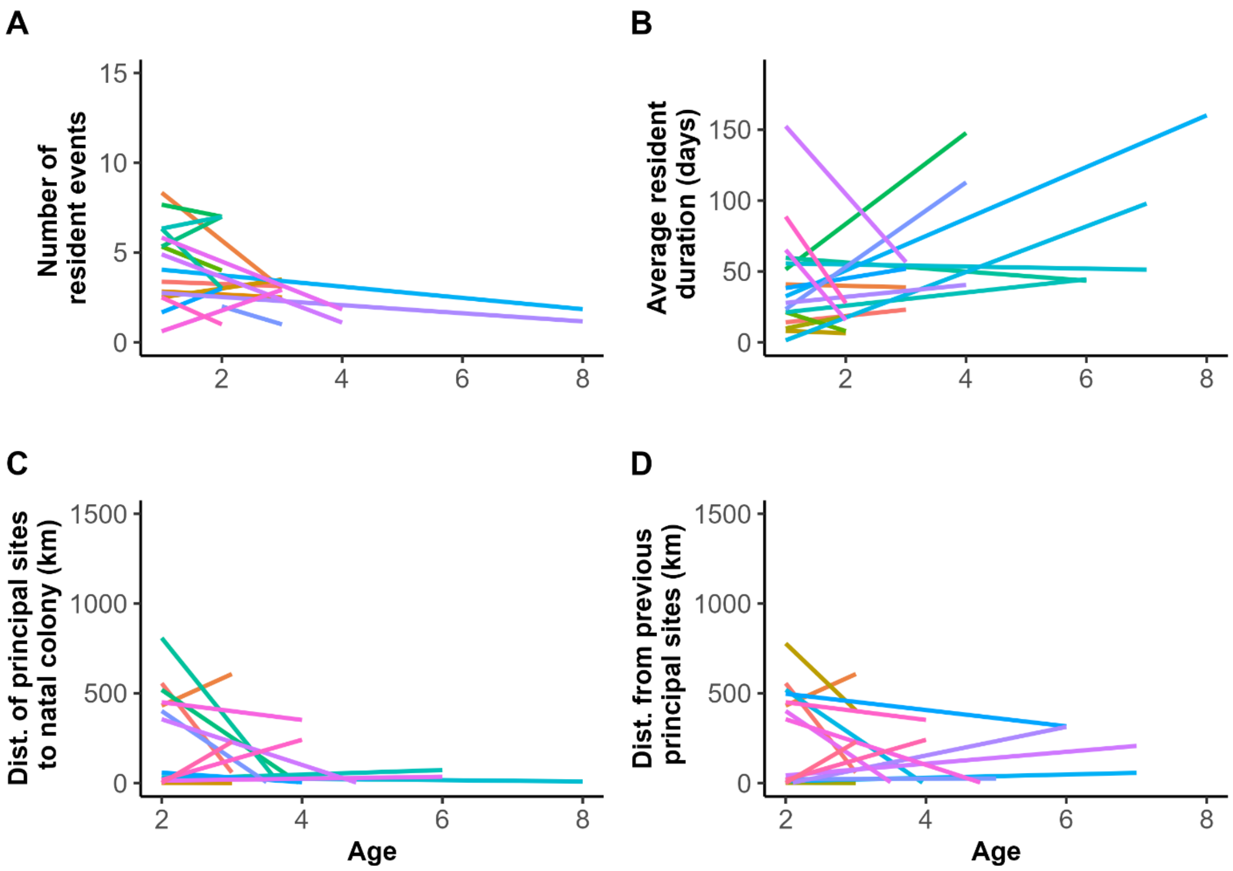


**Figure S5.** Individual-level ontogenetic shifts in residency metrics. A) number of staging events, B) average staging duration in days, C) distance of principal sites to natal the colony (km) and D) distance from previous principal sites (km). Results are shown for 25 individuals selected randomly.
